# Computational and experimental performance of CRISPR homing gene drive strategies with multiplexed gRNAs

**DOI:** 10.1101/679902

**Authors:** Samuel E. Champer, Suh Yeon Oh, Chen Liu, Zhaoxin Wen, Andrew G. Clark, Philipp W. Messer, Jackson Champer

## Abstract

CRISPR homing gene drives potentially have the capacity for large-scale population modification or suppression. However, resistance alleles formed by the drives can prevent them from successfully spreading. Such alleles have been found to form at high rates in most studies, including those in both insects and mammals. One possible solution to this issue is the use of multiple guide RNAs (gRNAs), thus allowing cleavage by the drive even if resistance sequences are present at some of the gRNA target sequences. Here, we develop a high-fidelity model incorporating several factors affecting the performance of drives with multiple gRNAs, including timing of cleavage, reduction in homology-directed repair efficiency due to imperfect homology around the cleavage site, Cas9 activity saturation, variance in the activity level of individual gRNAs, and formation of resistance alleles due to incomplete homology-directed repair. We parameterize the model using data from homing drive experiments designed to investigate these factors and then use it to analyze several types of homing gene drives. We find that each type of drive has an optimal number of gRNAs, usually between two and eight, dependent on drive type and performance parameters. Our model indicates that utilization of multiple gRNAs is insufficient for construction of successful gene drives, but that it provides a critical boost to drive efficiency when combined with other strategies for population modification or suppression.

## INTRODUCTION

An efficient gene drive could rapidly modify or suppress target populations^1–7^. These engineered constructs could potentially be used to prevent vector-borne diseases such as malaria or dengue and also have conservation applications^1–4^. The best studied form of gene drive is the homing drive, which utilizes the CRISPR/Cas9 system to cleave a wild-type allele. The drive is then copied into the wild-type site via homology-directed repair, increasing the frequency of the drive allele in the population. Thus far, CRISPR homing gene drives have been demonstrated in yeast^5–8^, flies^9–16^, mosquitoes^17–19^, and mice^20^.

However, homing drives suffer from high rates of resistance allele formation. Resistance alleles usually form when DNA is repaired by end-joining, which often results in a mutation of the sequence. This prevents targeting by the guide RNA (gRNA) and thus blocks subsequent conversion to a drive allele. Resistance alleles have been observed to form both in germline cells as an alternative to homology-directed repair and in the early embryo if a drive-carrying mother deposited Cas9 and gRNA into the egg^12^. While formation of resistance alleles remains the primary obstacle to construction of efficient gene drives, substantial progress has been made toward overcoming this challenge. For example, a suppression type drive in *Anopheles gambiae*^21^ and a modification type drive in *Drosophila melanogaster*^16^ avoided issues with resistance alleles by targeting an essential gene. Because of this, resistance alleles that disrupted the function of the target gene had substantially lower fitness than the drive. This allowed both drives to successfully spread through cage populations.

Multiplexing gRNAs has been proposed as a mechanism for increasing the efficiency of gene drives^1,4^. This would purportedly work by two mechanisms. First, having multiple cut sites would potentially allow drive conversion even if some of the sites have resistance sequences due to previous end-joining repair at those sites. As long as at least one site remains wild-type and thus, cleavable, homology-directed repair can still occur. Second, the chance of forming a full resistance allele that preserves the function of the target gene is substantially reduced due to possible disruptions at multiple gRNA target sites. Resistance alleles that disrupt the function of the target gene incur large fitness costs in several drive designs, which would make resistance substantially less likely to block the spread of the drive.

However, two studies utilizing two gRNAs^13,16^ showed somewhat lower increases in efficiency than that predicted by simple models of multiple gRNAs^22–24^. This is partially because most models assume that cleavage and repair by either homology-directed repair or end-joining occur sequentially at each gRNA target site. However, it appears that some resistance alleles in the germline form before the narrow temporal window for homology-directed repair^10,12,13^. Some may form as a direct alternative to homology-directed repair, but others appear to form after meiosis I when only end-joining repair is possible. Furthermore, unless cleavage occurs in both of the outermost gRNA target sites, the wild-type chromosome on either side of the cleavage would have imperfectly homology to the drive allele because of excess DNA between the cut and the homology arm ^13^. Imperfect homology likely reduces the fidelity of homology-directed repair and results in more end-joining repair. This proposition is supported by the greatly reduced efficiency seen in a construct with four gRNA targets far apart from one another^9^. Finally, it is unlikely that gRNAs are the limiting factor in Cas9/gRNA enzymatic activity^15^. As the number of gRNAs increases, we posit that the total cleavage rate plateaus, thus reducing the cleavage rate at each individual site and preventing further gains in drive efficiency.

Here, we systematically model these factors and show how they affect the performance of homing drives with multiple gRNAs. We confirm and parameterize these models via experimental analysis of several homing drives in *D. melanogaster*. We additionally consider other factors that could reduce gene drive performance, such as partial homology-directed repair and uneven activity of gRNAs. We then apply our model to drive performance in *Anopheles* mosquitoes, assessing several types of homing drives for population modification or suppression. We find that each type of drive has an optimal number of gRNAs that results in maximized overall performance, which could inform future designs of homing gene drives.

## RESULTS

### Simple model

To compare our results to previous work, we constructed a simple model of homing drive dynamics. This model considers each gRNA site completely independently with values inspired by highly efficient homing drives in *Anopheles* mosquitoes^18,19,21,25^. At each gRNA target site, there is a cut rate of 99%. If the site is cut, there is a 7.8% chance that a resistance sequence will be formed. Otherwise, homology-directed repair occurs, and the entire allele (including all target sites, even if some have resistance sequences) is converted to a drive allele. In this model, increasing the number of gRNA target sites increases the efficiency of the drive without limit (Figure 1). Even a few gRNAs are sufficient to reduce resistance allele formation to negligible levels. Under this model, multiple gRNAs were originally considered as a straightforward method to avert resistance in homing gene drives^22–24^.

**Figure 1.**
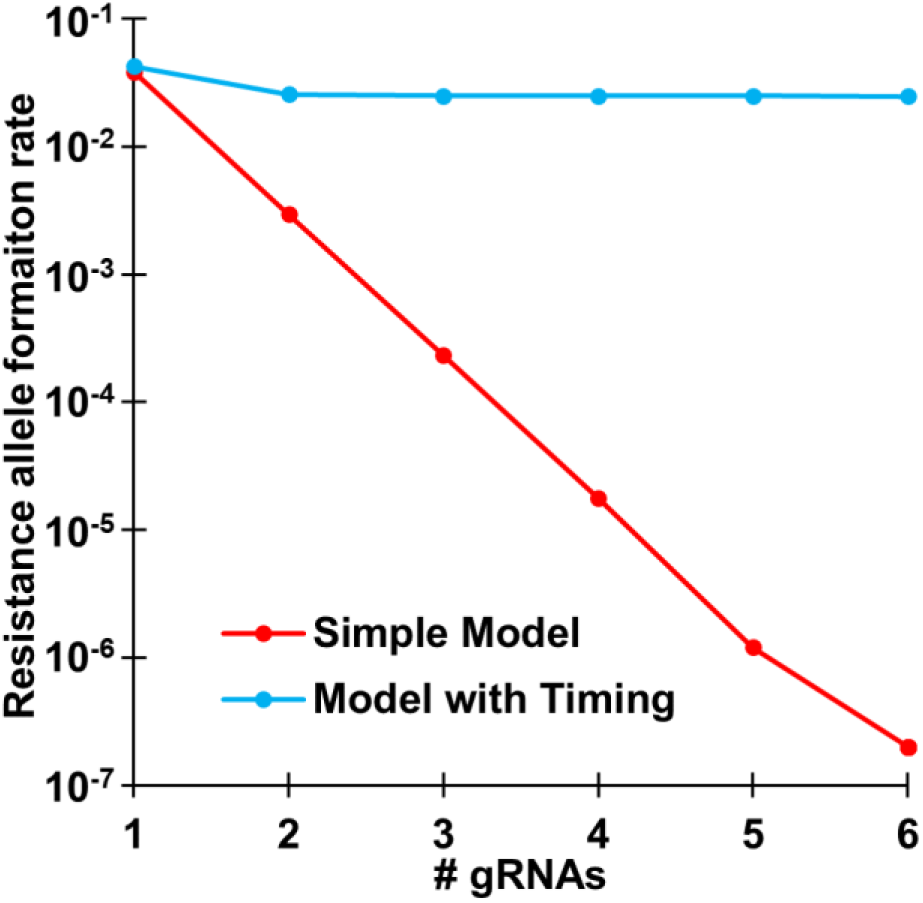
Resistance allele formation. Five million offspring were generated from crosses between drive/wild-type heterozygotes and wild-type individuals for each model and number of gRNAs. The rate at which wild-type alleles are converted to resistance alleles in the germline of drive/wild-type heterozygous individuals is shown.

### Model with timing

The simple model does not take into account timing of cleavage and homology-directed repair. Earlier experiments indicated that wild-type alleles can only be converted to resistance alleles and not to drive alleles in the early embryo due to maternally deposited Cas9^12,13^. Homology-directed repair of a chromosome does not take place at appreciable rates during this stage for the purposes of drive conversion. Furthermore, at least some resistance alleles that form in the germline do so in pre-gonial germline cells that can affect the genotype of multiple offspring^10,12,13^. After the chromosomes separate later in meiosis, homology-directed repair would no longer be possible, and any cleavage would result in formation of resistance alleles by end-joining repair. It is thus likely that there is only a narrow temporal window in the germline during which the drive can be successfully copied via homology-directed repair. This window likely covers early meiosis when homologous chromosomes are close together, which would increase the chance that one chromosome could be used as a template for repair of a double-strand break in the other. Thus, we constructed a model where cuts during a homology-directed repair phase occur simultaneously and have a single opportunity to undergo homology-directed repair. The model predicts that resistance alleles will form at or above a minimum baseline value equal to the chance that end-joining takes place during this phase instead of homology-directed repair (Figure 1). Additional gRNAs allow resistance to be reduced to close to this value, but not below it. Thus, the simple model may be inadequate to assess homing drive dynamics.

Previous experiments with two gRNAs indicated a lower efficiency improvement than even that predicted by our improved model that takes timing into account^13^. This was shown even more starkly with a four-gRNA drive^9^ that had a lower drive conversion efficiency than one-gRNA drives. We hypothesize that two additional factors account for this discrepancy. First, the rate at which homology-directed repair occurs after cleavage in the appropriate phase (which we refer to as repair fidelity) is certainly reduced if the DNA on either side of the cut sites doesn’t have immediate homology to the drive. Under these circumstances, the drive does not have DNA homologous to that found between the two outermost cut sites. This tends to reduce the efficiency of the drive unless both outer gRNAs are cleaved. Second, the amount of Cas9 enzyme is limited, so as the number of gRNAs increases, Cas9 becomes saturated with gRNAs and cleavage activity plateaus. This has the effect of decreasing the cleavage rate at individual gRNA sites as the total number of gRNAs increases. To test the impact of repair fidelity and Cas9 activity saturation, we conducted a series of experiments.

### Synthetic target site experiments with one gRNA

We first constructed a one-gRNA system in *D. melanogaster* that targeted EGFP with a drive containing dsRed and with a Cas9 gene driven by the *nanos* promoter (Figure S1), similar to previously demonstrated synthetic target site drives^15^. Drive/wild-type heterozygotes displayed a drive conversion efficiency of 83% in females and 61% in males (Data S1). These values were higher than previous synthetic target site drives^15^, likely due to the difference in genomic location of the target site or the gRNA, which targeted further away from the 3xP3 promoter in EGFP.

Since multiplexing of gRNAs can best be accomplished by expressing them from a single compact promoter, we constructed a one-gRNA system identical to the above, but with a tRNA that must be spliced out of the gRNA gene to yield an active gRNA. By including additional tRNAs between gRNAs, several gRNAs can be expressed together with this system^26^. We found that drive/wild-type heterozygote females had a drive conversion efficiency of 82% in females and 65% in males (Data S2). This indicates that the tRNA system is fully capable of supporting efficient gene drives.

We next constructed a drive to determine the effects of poor homology between the cleaved wild-type chromosome and the drive allele. To accomplish this, we used a single gRNA as above with the tRNA system, but we aligned the right homology arm to a hypothetical second gRNA cut site, instead of to the first cut site (Figure 2). Thus, the first 114 nucleotides on the right arm of the cut site would not be homologous to any DNA around the drive allele. Drive conversion rates for females were only 84% of the rate of the one-gRNA drive that had full homology around the cut site, while the rate for males was 89% of that of the full homology drive (Data S3). This indicates that a multiple-gRNA drive would indeed exhibit lower conversion efficiency when cleavage does not take place at both ends.

**Figure 2.**
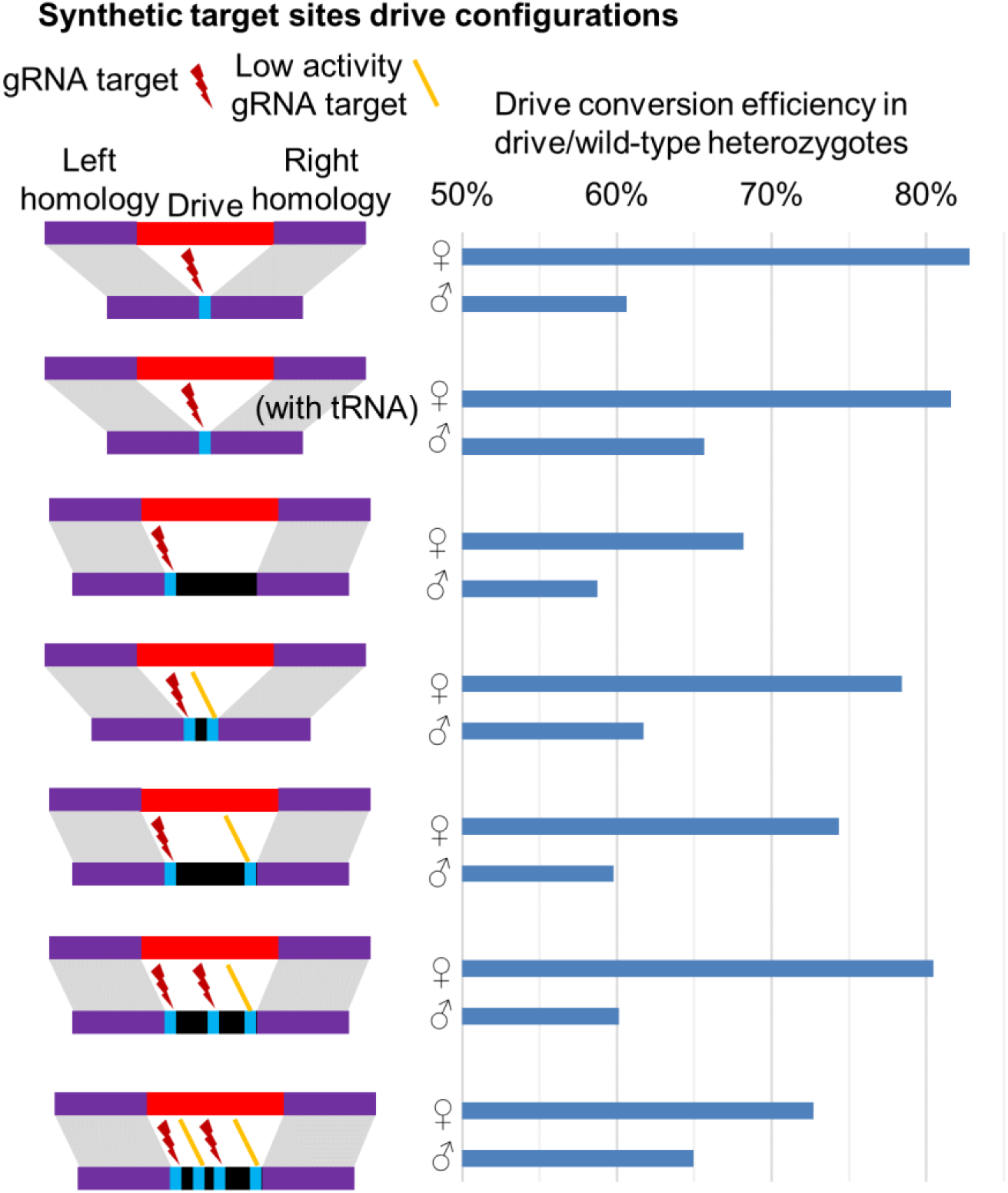
Experimental performance of homing drives with different configurations. Blue shows the gRNA target sites and black shows regions of DNA that have no homology to the drive alleles. Note that active gRNAs are shown by a dark red lightning bolt, and gRNAs with very low activity are shown with an orange line icon.

### Split drive experiments

To assess the effects of saturation, we examined three constructs containing Cas9 with either zero, one, or four gRNAs targeting a genomic region between two genes and downstream of both. Mutations from cleavage in this area are thereby unlikely to affect an individual’s fitness or other characteristics. These constructs were placed at the same genomic site as the one-gRNA synthetic target site constructs in this study. Individuals with these constructs were crossed to the split-drive targeting *yellow* that we developed previously^15^ to generate individuals heterozygous for a Cas9 element and a split-drive element. The embryo resistance allele formation rates in individuals with zero, one, or four gRNAs in the Cas9 element were 83%, 72%, and 65%, respectively (Data S4). The amount of the gRNA targeting *yellow* was constant in these drives, and increasing quantities of other gRNAs decreased the rate at which *yellow* was cleaved. This is consistent with the hypothesis that saturation of Cas9 activity reduces the cleavage rates of drives with multiple gRNAs.

Nevertheless, Cas9 does not necessarily become fully saturated with a single gRNA. The total cleavage rate could potentially somewhat increase if additional gRNAs are provided, even though it would likely plateau rapidly. When heterozygotes for the split drive targeting *yellow*^15^ and the standard drive targeting *yellow*^12^ with one copy of Cas9 and two gRNA genes were crossed to *w*^*1118*^ males, the rate of embryo resistance allele formation and mosaicism was somewhat higher than for standard drive/resistance allele heterozygotes with one copy of Cas9 and only one gRNA gene (Data S5).

### Experiments with multiple gRNAs

To assess the performance of drives with multiple gRNAs, we constructed several additional constructs targeting EGFP, but with two, three, or four gRNAs (Figure 3). The left target site for each of these was the same as the one-gRNA synthetic target site drives, and the homologous ends of all of these drives match the left and right gRNA target sites. However, we found that of the four gRNAs used, only the first and the third had substantial cleavage activity, as indicated by sequencing of embryo resistance alleles (Table S1). Though germline cleavage activity was likely somewhat higher than in the embryo for these gRNAs, their low activity undoubtedly reduced drive performance. Nevertheless, we found that the overall performance of these drives was consistent with our predictions of the effects of timing, repair fidelity, and Cas9 activity saturation (Figure 3). The results clearly show that adding additional gRNAs does not result in highly efficient drives.

**Figure 3.**
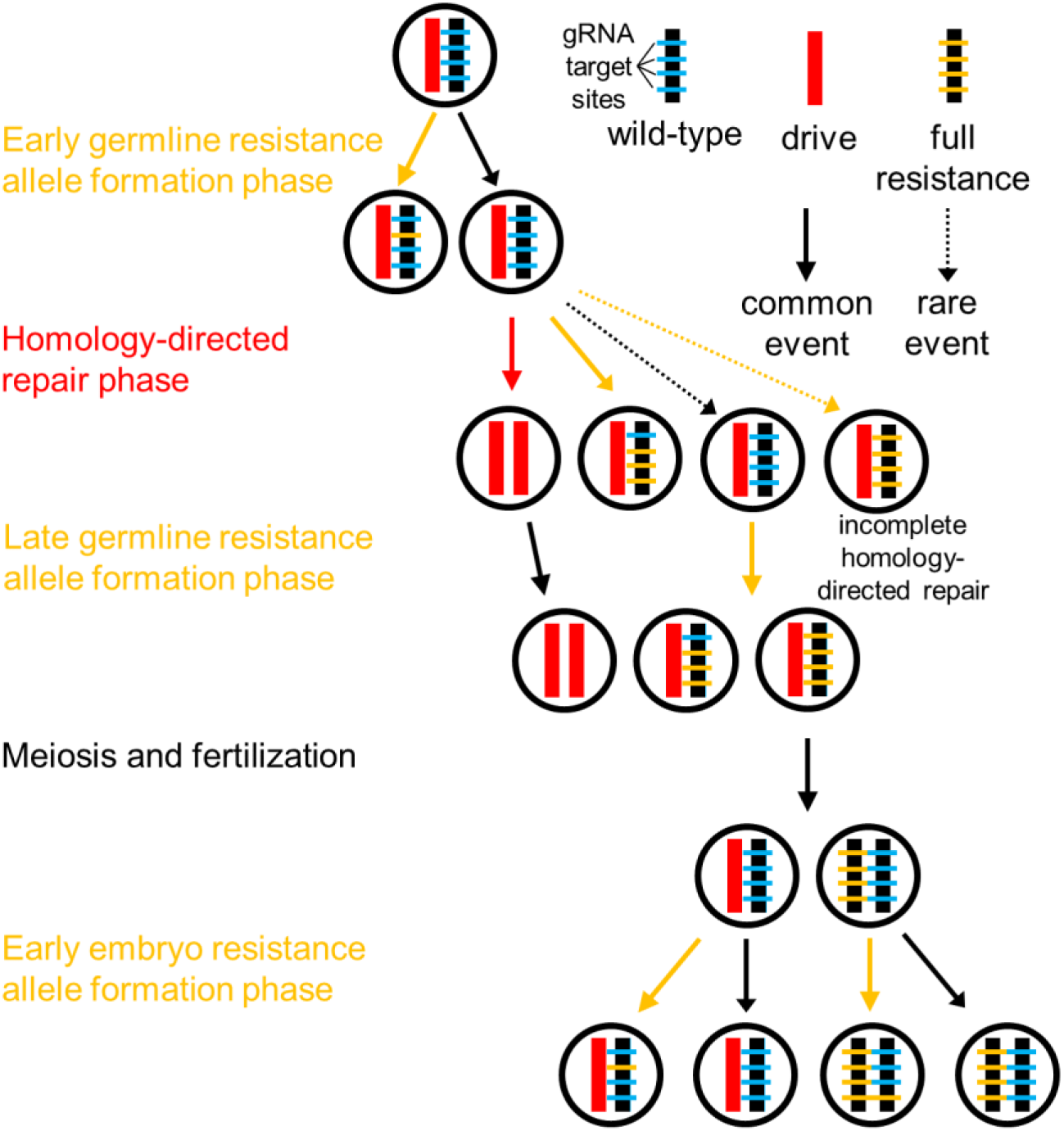
Steps in the model. First, wild-type gRNA target sites can be cleaved in the early germline, forming resistance alleles. During this process, simultaneous deletions can occur, and individual resistance sequences can either disrupt or preserve the target gene function. Next, cleavage occurs at a high rate in the homology-directed repair phase. Usually, this results in successful conversion to a drive allele. However, if homology-directed repair fails to occur, end-joining can form resistance alleles. Incomplete homology-directed repair can also convert the entire allele to a resistance allele, ignoring individual target sites. Next, another resistance allele formation phase converts most remaining wild-type sites into resistance sequences. Meiosis and fertilization take place, and then, if the female parent had at least one drive allele, a final round of resistance allele formation takes place in the early embryo.

Specifically, we constructed two two-gRNA drives. One of these had two closely spaced gRNA targets (36 nucleotides apart) and showed a drive conversion efficiency of 78% in females and 62% in males (Data S6). This was higher than the two-gRNA drive with gRNAs that were more widely spaced (114 nucleotides apart), which demonstrated drive conversion efficiencies of 74% in females and 60% in males (Data S7). With the second gRNA having low activity in each of these drives, the difference between them is accounted for by the lower repair fidelity in the drive with more widely spaced gRNAs (Figure 3). The drive with three gRNAs was similar to the two-gRNA drive with widely spaced gRNAs, with the addition of a third active gRNA in between them. This increases the overall cleavage rates due to the higher proportion of active gRNAs and allows for greater repair fidelity on the right end, since cleavage in this system usually takes place at the left and middle gRNA targets, instead of only at the left gRNA target. Thus, this construct showed an improved drive conversion efficiency of 80% in females, though male drive conversion efficiency apparently remained at 60%. A final construct added an additional gRNA between the left and middle gRNAs (the same gRNA that the closely spaced two-gRNA construct added). However, since this gRNA had low activity, overall drive performance was negatively affected by saturation of Cas9 by gRNAs, resulting in a reduced drive conversion efficiency of 73% in females, though male drive conversion efficiency appears to have improved to 65% (possibly due to an underestimation in the 3-gRNA construct).

### Refinement of the model

To refine our model, we incorporated distinct phases to fully account for homing drive dynamics in the germline (Figure 3). First, early germline resistance alleles form, followed by a homology directed repair phase and a late germline resistance allele formation phase. In the embryos of mothers with at least one drive allele, maternally deposited Cas9 and gRNA can result in the formation of additional resistance alleles.

We additionally model reduced repair fidelity from lower homology around the cut sites, Cas9 activity saturation, and variance in the activity level of individual gRNAs. See the Supplemental Results section for a detailed treatment of these model components and estimation of parameters based on our experiments.

Models with repair fidelity or Cas9 activity saturation alone did not produce much deviance from our basic model with timing (Figure 4). However, in a model that includes both repair fidelity and Cas9 activity saturation, we find the emergence of an optimal number of gRNAs to maximize drive conversion efficiency (Figure 4), with drive conversion decreasing rapidly after this optimal level. gRNA activity variance has only a small effect on drive conversion performance (Figure 4). With our default parameters simulating an efficient *A. gambiae* construct, the optimal number of gRNAs is two, though drives with three gRNAs have nearly as good conversion efficiency. However, the optimal number of gRNAs for a drive may be greater than the optimal number for drive conversion efficiency, as detailed below.

**Figure 4.**
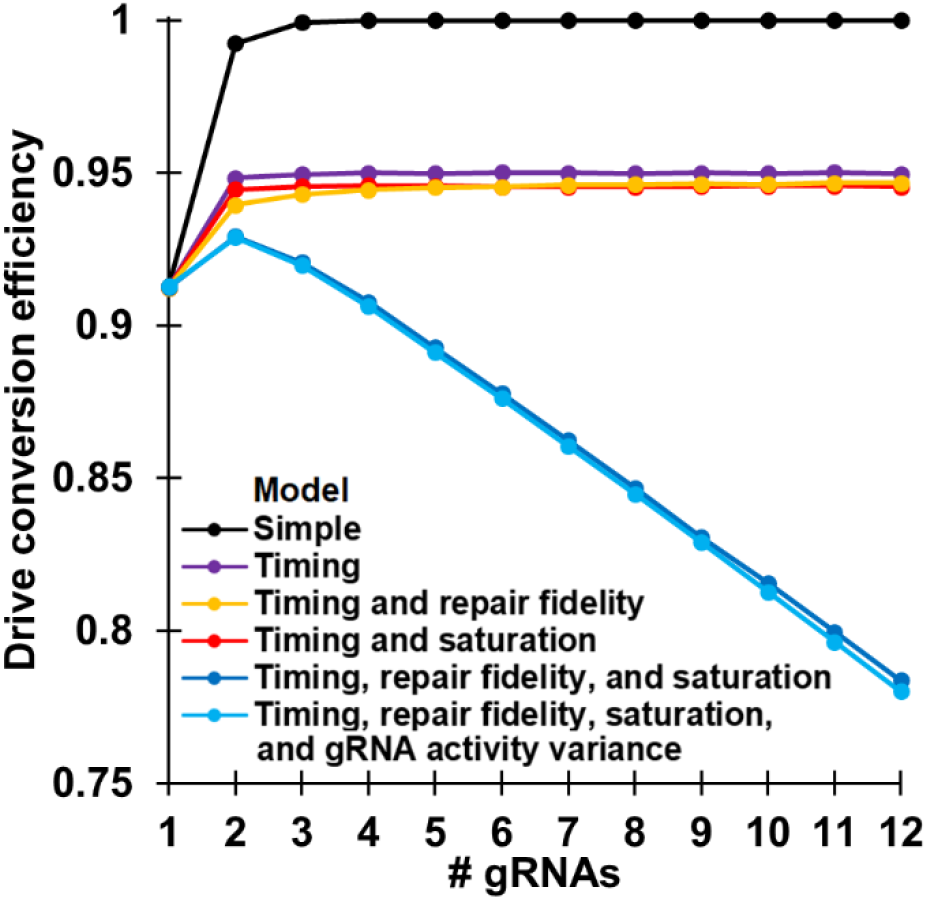
Effects of model components on drive performance. Five million offspring were generated from crosses between drive/wild-type heterozygotes and wild-type individuals for each model and number of gRNAs. The rate at which wild-type alleles are converted to drive alleles in the germline of drive/wild-type individuals is shown.

### Types of resistance alleles

Resistance alleles can be divided into two classes. The first class, which preserves the function of any target gene, is rarer. Resistance alleles that disrupt the target gene are more common due to frameshift mutations or other disruptions to the target sequence. In our model, we assume no special targeting of conserved sequences and thus estimate that resistance sequences that preserve the function of the target gene form in 10% of cases^12,13^. In reality, this could be substantially reduced^13,16,21^. In our model, we further assume that a resistance sequence preserving the function of the target gene must form independently at each gRNA target site for the target gene to retain its function. If even a single resistance sequence that disrupts the function of the target gene is present, the target gene is assumed to be non-functional. Any deletion due to simultaneous cleavage followed by end-joining repair is also assumed to disrupt the target gene. Thus, one major advantage of multiple-gRNA drives is that complete resistance alleles that preserve the function of the target gene become exponentially less common as the number of gRNAs increases (Figure 5, black line).

**Figure 5.**
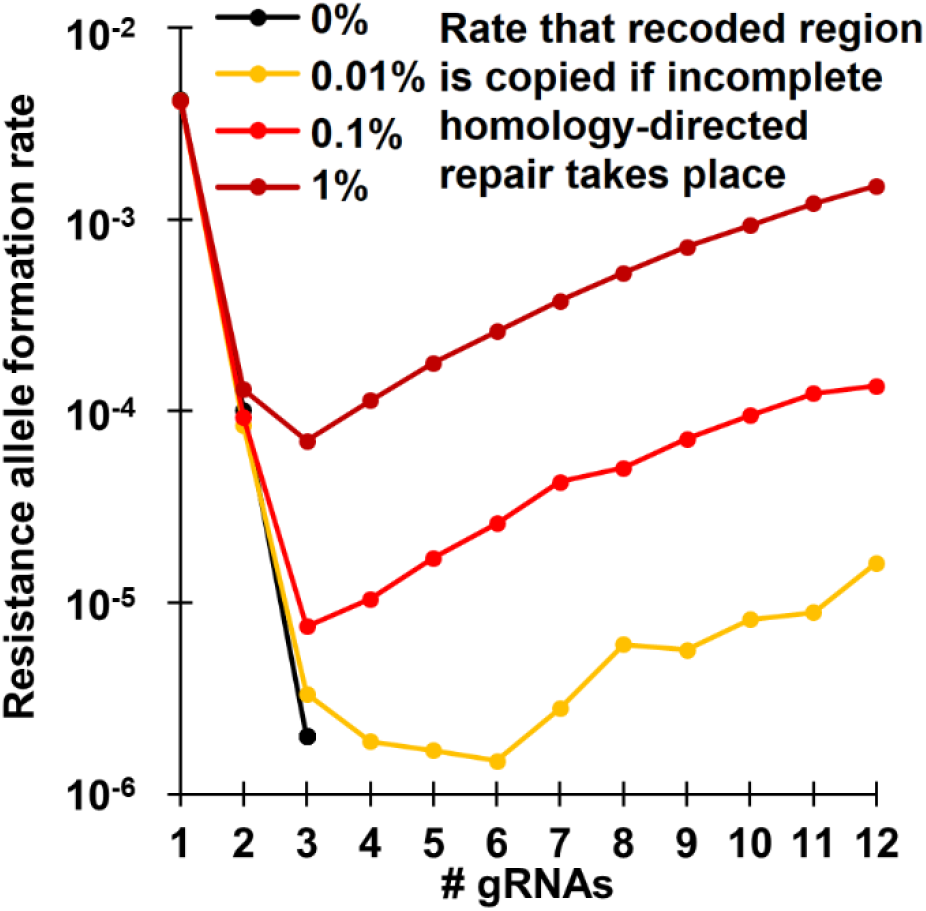
Formation of resistance alleles that preserve the function of the target gene. Five million offspring were generated from crosses between drive/wild-type heterozygotes and wild-type individuals using the full model for each number of gRNAs and value of the parameter determining the chance that incomplete homology-directed repair results in the formation of resistance alleles that preserve the function of the target gene. That rate at which resistance allele that preserve the function of the target gene is shown. Note that no such resistance alleles were formed when incomplete homology-directed repair was incapable of forming these alleles and if at least four gRNAs were present.

However, gene drives containing a recoded sequence of the target gene that is immune to cleavage are vulnerable to another mechanism for forming resistance alleles that preserve the function of the target gene. If homology-directed repair copies the recoded portion of the drive without copying the drive’s payload, such a resistance allele will form regardless of the number of gRNAs. This is further described in the Supplemental Results section covering incomplete homology-directed repair. This results in an optimal number of gRNAs in such drives for minimization of resistance alleles that preserve the function of the target gene (Figure 5).

### Results of the full model for multiplexed gRNAs

With our model in place, we consider the performance of several types of drives. The first of these is the standard homing drive, in which resistance allele classes have no particular effect. The next drive is for population suppression^21^, which is conducted by targeting a recessive female fertility gene. For this drive, females are rendered sterile unless they possess at least one wild-type allele or resistance allele that preserves the function of the target gene. We also consider approaches for population modification that target a haplolethal or recessive lethal gene, where the drive has a recoded sequence of the gene that cannot be cleaved by the drive’s gRNAs^16^. In the haplolethal approach, any individual with a resistance allele that disrupts the target gene is nonviable, removing these alleles from the population. In the recessive lethal approach, an individual is only nonviable if it has two such resistance alleles. Finally, we consider a population modification approach that targets a gene of interest, such as a gene required for malaria transmission in *Anopheles*^27,28^. Rather than carrying a payload, this drive’s purpose is to disrupt its target in a manner similar to that of the suppression drive.

We found that the optimal number of gRNAs in the population modification drives to achieve a maximum drive frequency was three, though drives with two gRNA were nearly as efficient (Figure 6A). Due to their ability to remove resistance alleles quickly, the haplolethal drive reached nearly 100% frequency (Figure 6A). The recessive lethal drive is slower at removing resistance alleles when they are rare, so it reached a lower frequency (Figure 6A). However, the haplolethal drive removes drive alleles when they are present in the same individual as a resistance allele that disrupts the function of the target gene. Thus, this system spreads somewhat more slowly than other types of population modification drives, though not as slowly as the population suppression homing drive (Figure 6B). Of particular interest, gRNAs beyond two reduce drive conversion efficiency, which results in a slower spread of the drive (Figure 6B). However, having multiple gRNAs is essential for reducing the formation rate of resistance alleles that preserve the function of the target gene (Figure 6C). For homing drives with payloads (standard, haplolethal, and recessive lethal), the optimal number of gRNAs is three. Greater amounts slow down the drive and result in increased resistance allele formation for the haplolethal and recessive lethal drive due to incomplete homology-directed repair. Thus, having three gRNAs is usually optimum for maximizing drive frequency after 100 generations (Figure 6D). However, for the gene disruption homing drive, the optimal number of gRNAs for maximizing the frequency of “effector” alleles was 4-6 (Figure 6D). This is because effector alleles for this drive include not only the drive, but also resistance alleles that disrupt the function of the target gene. Together with the lack of formation of resistance alleles that preserve the function of the target sequence via incomplete homology-directed repair, this enables the gene disruption drive to make efficient use of a higher number of gRNAs. Drives with somewhat reduced performance modeled after our *Drosophila* experiments show similar patterns, although the optimal number of gRNAs can be slightly higher than models based on *Anopheles* drives (Figure S5, and see Supplemental Results).

**Figure 6.**
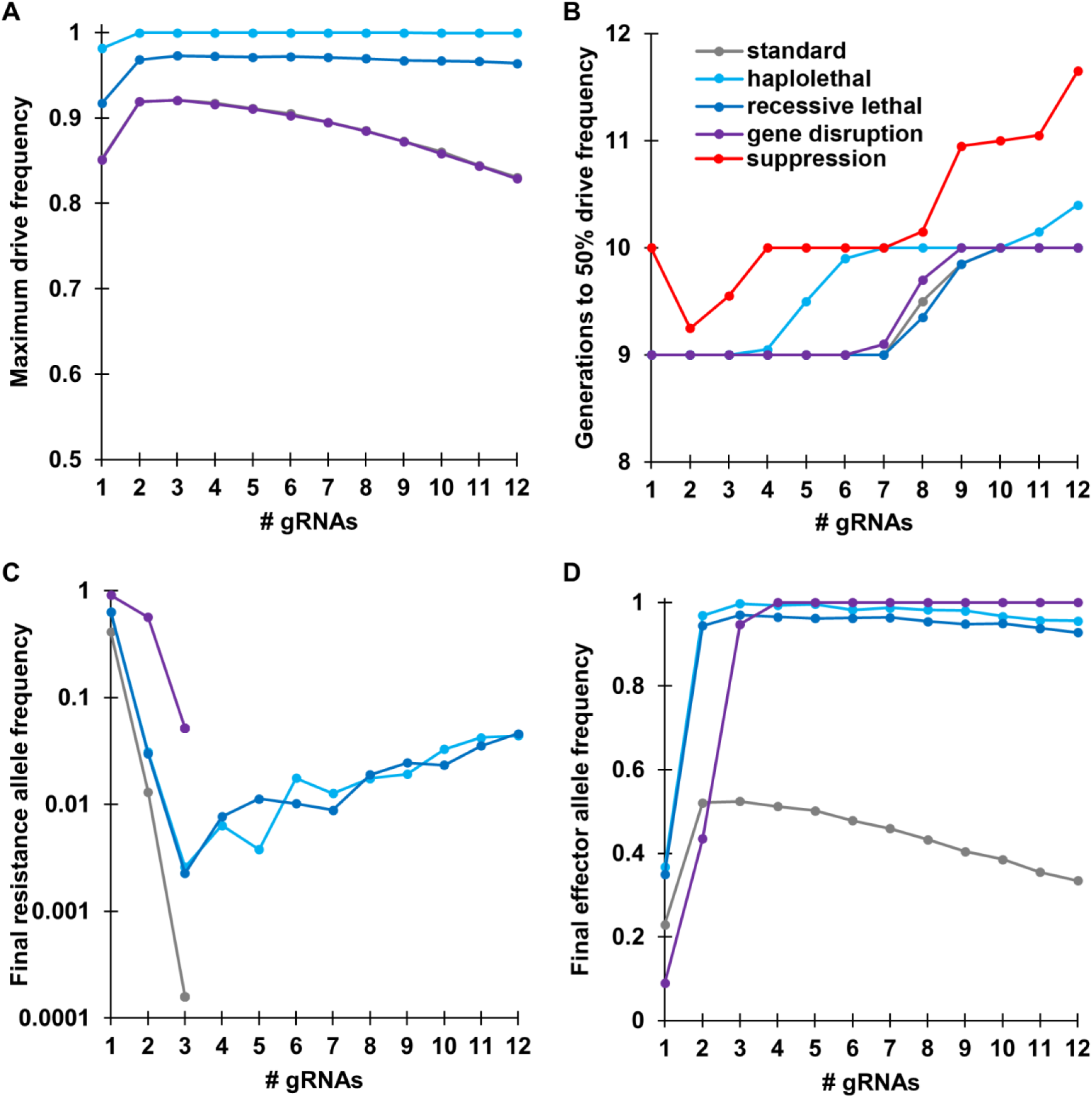
Comparison of performance parameters for different types of homing drive. Drive/wild-type heterozygotes were released into a population of 100,000 individuals at an initial frequency of 1%. The simulation was then conducted for 100 generations using the full model. The displayed results are the average from 20 simulations for each type of drive and number of gRNAs. (**A**) The maximum drive allele frequency reached at any time in the simulations. Note that the standard drive and gene disruption drive values are highly similar. (**B**) The number of generations needed for the drive to reach at least 50% total allele frequency. Note that the suppression drive is only shown in (**B**). (**C**) The final frequency of resistance alleles after 100 generations. The displayed values are only for resistance alleles that preserve the function of the target gene. No resistance alleles were present in the standard drive and gene disruption drive when at least four gRNAs were present. (**D**) The final effector frequency present in the population after 100 generations. This was the drive allele only for most drive types, but for the gene disruption drive, it includes resistance alleles that disrupt the function of the target gene.

### Multiplexing gRNAs for suppression drives

Suppression type homing drives are particularly sensitive to both the resistance allele formation rate and the rate that the drive spreads through a population. When examining the rate of successful suppression (Figure 7), our high-performance drives with default parameters were successful at suppressing the population if there were a sufficient number of gRNAs to avoid formation of resistance alleles that preserve the function of the target gene. However, a drive with somewhat reduced performance (see supplemental results) was unable to always achieve successful suppression, regardless of the number of gRNAs. Furthermore, when the number of gRNAs was greater than eight, the rate of successful suppression declined rapidly due to reduced drive conversion efficiency (Figure 7), resulting in an optimum number of gRNAs between four and eight.

**Figure 7.**
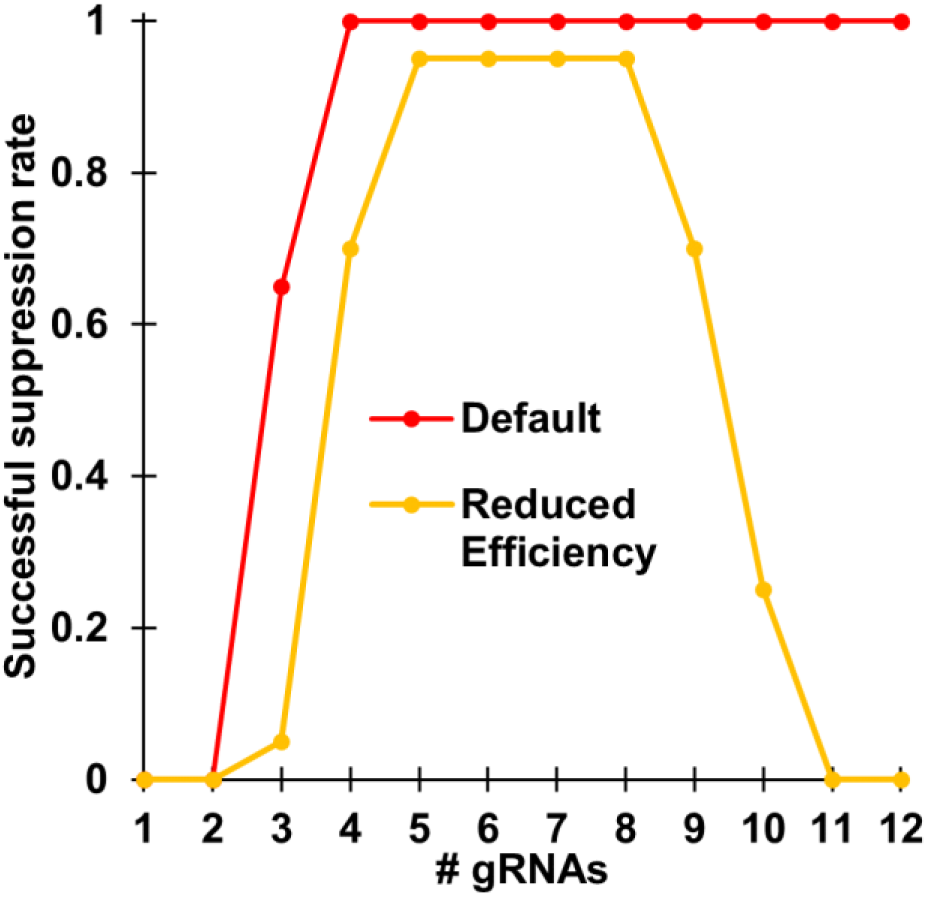
Number of gRNAs needed for successful population suppression. Drive/wild-type heterozygotes with a suppression drive were released into a population of 100,000 individuals at an initial frequency of 1%. The simulation was then conducted for 100 generations. The displayed results are the average from 20 simulations for each type of drive and number of gRNAs. The fraction of simulations that resulted in successful suppression are shown. The full model was used, which included an early germline cleavage rate of 2%, a homology-directed repair phase cleavage rate of 98%, and an embryo cleavage rate of 5%. For the reduced efficiency model, these parameters were changed to 5%, 92%, and 10%.

## DISCUSSION

Based on our results, homing drives likely have an optimal number of gRNAs that maximize drive efficiency while reducing the formation rate of resistance alleles that preserve the function of the target gene to an acceptable level. This result emerged naturally from a model with specific time steps, Cas9 activity saturation, and reduced repair fidelity when homology ends around the cuts sites fail to line up perfectly. Even with a more basic model that includes a narrow window for homology-directed repair, we can reject the notion that having sufficiently high numbers of gRNAs results in highly efficient drives.

Overall, we show that while multiple gRNAs are useful for improving drive efficiency and reducing resistance, these performance gains are far smaller than that predicted by simple models with sequential cutting and repair^22–24^ or even models that allow for simultaneous cutting^22^. Earlier work indicated that the window for homology-directed repair is narrow, with only resistance alleles forming before and afterward^10,12,13^. A better understanding of this window, rates of successful homology-directed repair, and proportion of resistance alleles formed before, during, and after this window would allow for improvements to our model. Homology of DNA on either side of a cut site is well-known to be critical for the fidelity of homology-directed repair, and we showed that it indeed influences drive conversion efficiency. Finally, Cas9 cleavage activity certainly reaches a maximum as more gRNAs are added. Though it remains unclear when exactly this occurs, it is likely that for many gRNA promoters, a maximum cut rate would be quickly reached, thus reducing the cleavage rates of individual gRNAs as the total number is increased. Future studies could investigate how such saturation occurs and enable refinement of the quantitative model predicting activity.

Indeed, while our model represents a step forward in our understanding of how multiplexed gRNAs affect homing drive efficiency, further improvements are needed to be able to more accurately predict homing drive performance. In particular, the rate of resistance allele formation from incomplete homology-directed repair could be better quantified, with particular attention paid to the rate at which any recoded region is fully copied, forming a resistance allele that preserves the function of the target gene. Calculations on the rate of formation of such alleles from end-joining may also need to be revised. In our model, we assumed that each gRNA cut site independently had the same chance of forming a resistance sequence that disrupts the function of the target gene. In practice, frameshift mutations between gRNA cut sites, but with restored frame after the last mutated site, may be insufficient to disrupt the function of the target gene. Thus, a good practice to minimize formation of resistance alleles that preserve the function of the target gene may be to target conserved or important regions less tolerant of mutations, and perhaps to space gRNAs far enough apart, despite reduced drive conversion, to ensure that a frameshift between any two gRNA sites will disrupt the gene. At minimum, gRNAs should be placed be far enough apart to prevent mutations at one site from converting an adjacent target site into a resistance allele. Finally, variance in the activity level of gRNAs is well known, and we also observed this in our multiple gRNA homing drives in this study. Such activity levels could potentially be predicted^29^, but experimental assessment will likely remain necessary in the immediate future.

Our models are informative about the relative strengths and weaknesses of the different types of homing gene drives. Standard drives lack any particular mechanism for removing resistance alleles that disrupt the function of the target gene (indeed, they may not even target a specific gene), which means that a successful drive of this nature requires a high drive efficiency, very low resistance allele formation rates, and low fitness costs for to persist long enough to provide substantial benefits. The optimal number of gRNAs for such drives is likely low, perhaps two or three for a highly efficient system.

Drives that target haplolethal or recessive lethal genes can effectively remove resistance alleles that disrupt the function of the target gene, allowing such drives to tolerate substantially higher overall rates of resistance allele formation. These drives do not lose much efficiency with large numbers of gRNAs because even though drive conversion efficiency is reduced, the drives also operate by toxin-antidote principles^30,31^, enabling removal of wild-type alleles and an accompanying relative increase of drive allele frequency even without drive conversion. However, we hypothesize that with reduced homology around the cut sites, incomplete homology-directed repair becomes more likely, thus forming a balance between formation of resistance alleles that preserve the function of the target gene due to partial homology-directed repair and end-joining mechanisms. It is unclear how common the former phenomenon is, but it seems likely that the optimal number of gRNAs for such drives is perhaps three or four. This is mostly influenced by the rate of partial homology-directed repair, which could perhaps be minimized if the drive was located in an intron (which could be a synthetic intron not present in the wild-type genome) with essential recoded regions on either side of the intron, allowing for efficient use of a greater number of gRNAs. This may not be necessary, however, if the rate of resistance allele formation that preserves the target site is substantially less than the rate at which any payload gene is inactivated by mutations that occur during homology-directed repair (10^−6^/nucleotide), which is approximately 1,000-fold greater than the rate by DNA replication. If such a rate would preclude effective deployment of a homing drive, then toxin-antidote systems^30,31^ that rely only on DNA replication for copying of payload genes may be more suitable.

A gene disruption drive for population modification could potentially avoid both the need for a recoded region and inactivation of payload genes by targeting an endogenous gene. In this case, the end goal would be to disrupt this gene either by the presence of the drive or by formation of resistance alleles. These drives do not need a payload. In this case, formation of resistance alleles that disrupt the target gene may actually be beneficial due to their reduced fitness cost compared to the drive. For such a drive, the optimal number of gRNAs would be the minimum number necessary to prevent formation of resistance alleles that preserve the function of the target gene, perhaps four to eight, depending on population size and drive performance.

A suppression type drive has similar considerations, but with a narrower window for success. This is because any formation of resistance alleles that preserve the target gene would likely result in immediate failure of the drive. Additionally, if drive conversion efficiency is insufficient, the drive may lack the power to suppress the population in the first place, at least in a reasonable timeframe. Thus, a narrower range of four to six gRNAs is likely optimal for such a drive. For all of these drive types, if the rate of resistance allele formation that preserves the function of the target gene is lower than in our models (such as by targeting a sequence that is highly intolerant of mutations^21^), the optimal number of gRNAs will be somewhat reduced. Overall, we conclude that gRNAs sites should be placed as close together as possible, while still far enough apart to prevent mutations at one target site from affecting adjacent sites. The total number of gRNA should be kept relatively low: at least two, but well under a dozen, with the exact number depending on the type of drive and other performance characteristics. While multiplexing of gRNAs is unlikely to enable the success of a homing drive without a supporting strategy, it will likely be a critical component of a successful drive. Due to the simplicity of this strategy, we expect that it will be increasingly common, enabling refinement of models, which will in turn allow for the more rapid construction of efficient homing drives with less need for optimization.

## ACKNOWLEDGEMENTS

This study was supported by funding from New Zealand’s Predator Free 2050 program under award SS/05/01 to P.W.M. and the National Institutes of Health awards R01GM127418 to P.W.M., R21AI130635 to J.C., A.G.C., and P.W.M, and F32AI138476 to J.C.

## SUPPLEMENTAL INFORMATION

### METHODS

#### Plasmid construction

The starting plasmids pCFD3^32^ (Addgene plasmid #49410) and pCFD5^26^ (Addgene plasmid #73914) were kindly supplied by Simon Bullock, and starting plasmids IHDyi2^12^, and BHDgN1a^15^, and p3xP3-dsRedv2^15^ were constructed in our previous studies. Plasmid digests were conducted with restriction enzymes from New England Biolabs (HF versions, when possible). PCR was performed with Q5 Hot Start DNA Polymerase (New England Biolabs), and DNA oligos and gBlocks were obtained from Integrated DNA Technologies. Gibson assembly of plasmids utilized Assembly Master Mix (New England Biolabs), and plasmids were transformed into JM109 competent cells (Zymo Research). Plasmids used for injections were purified using the ZymoPure Midiprep kit (Zymo Research). Cas9 gRNA target sequences were found with CRISPR Optimal Target Finder^33^. The following tables show the DNA fragments used for Gibson Assembly of each plasmid.

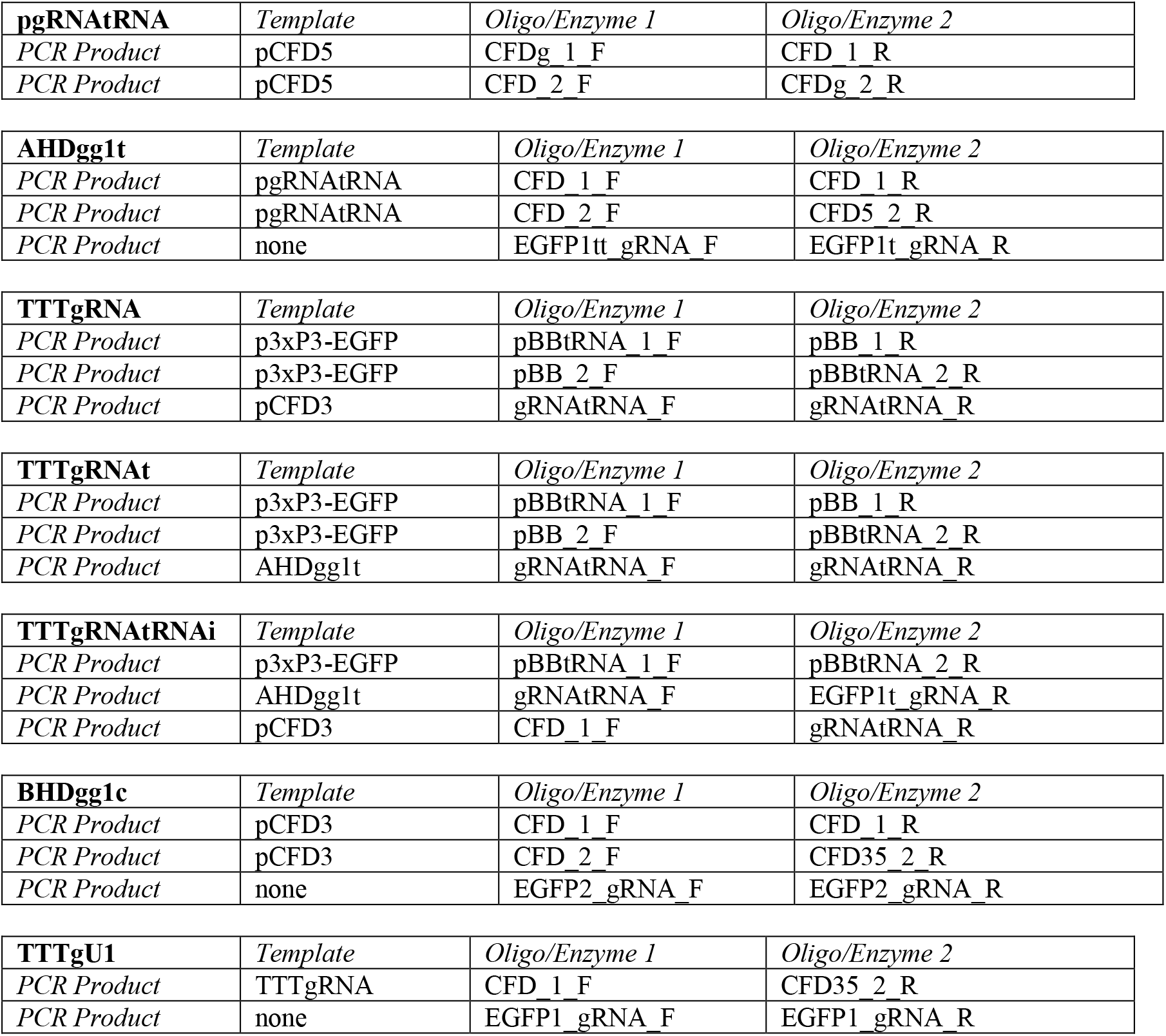
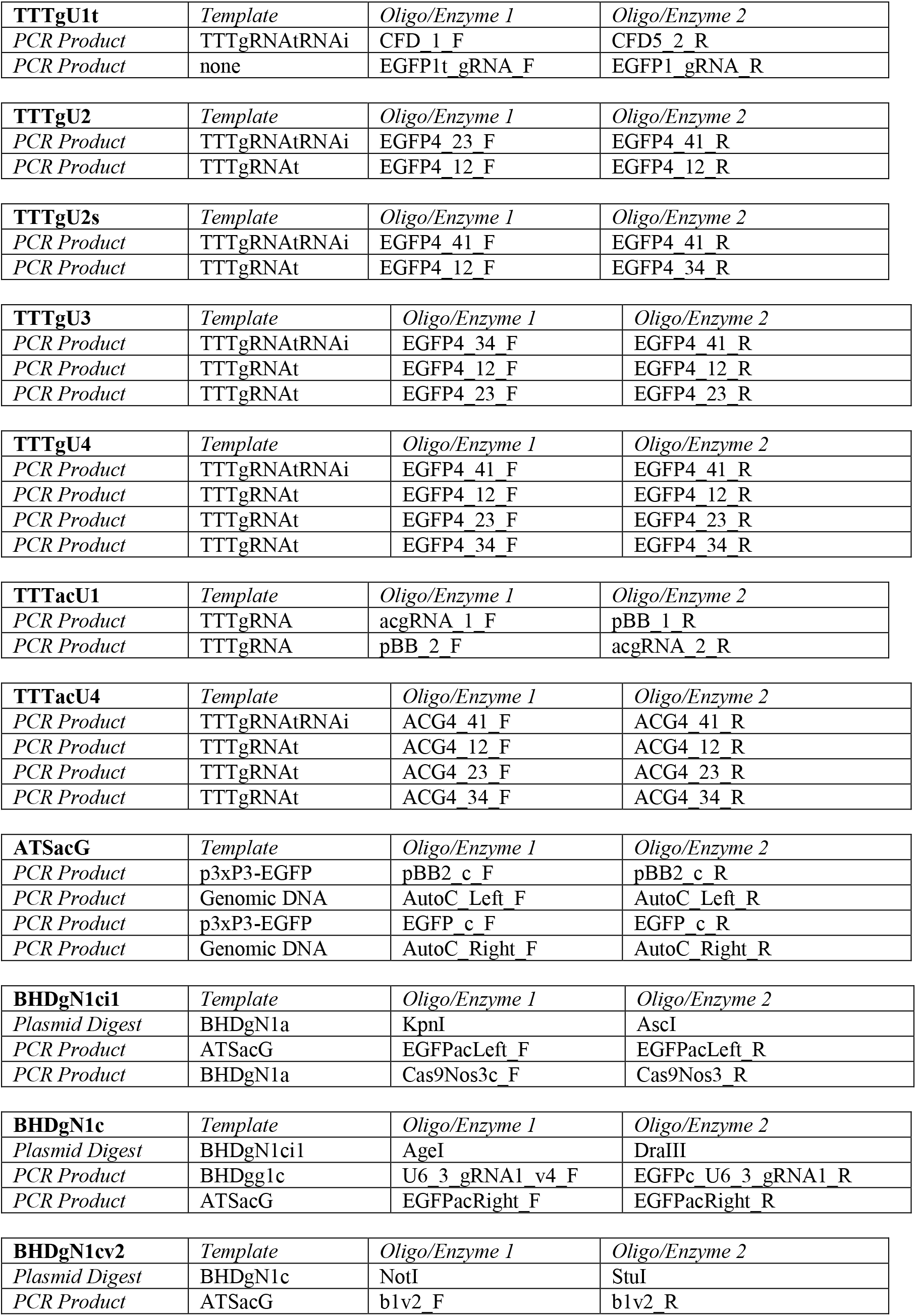
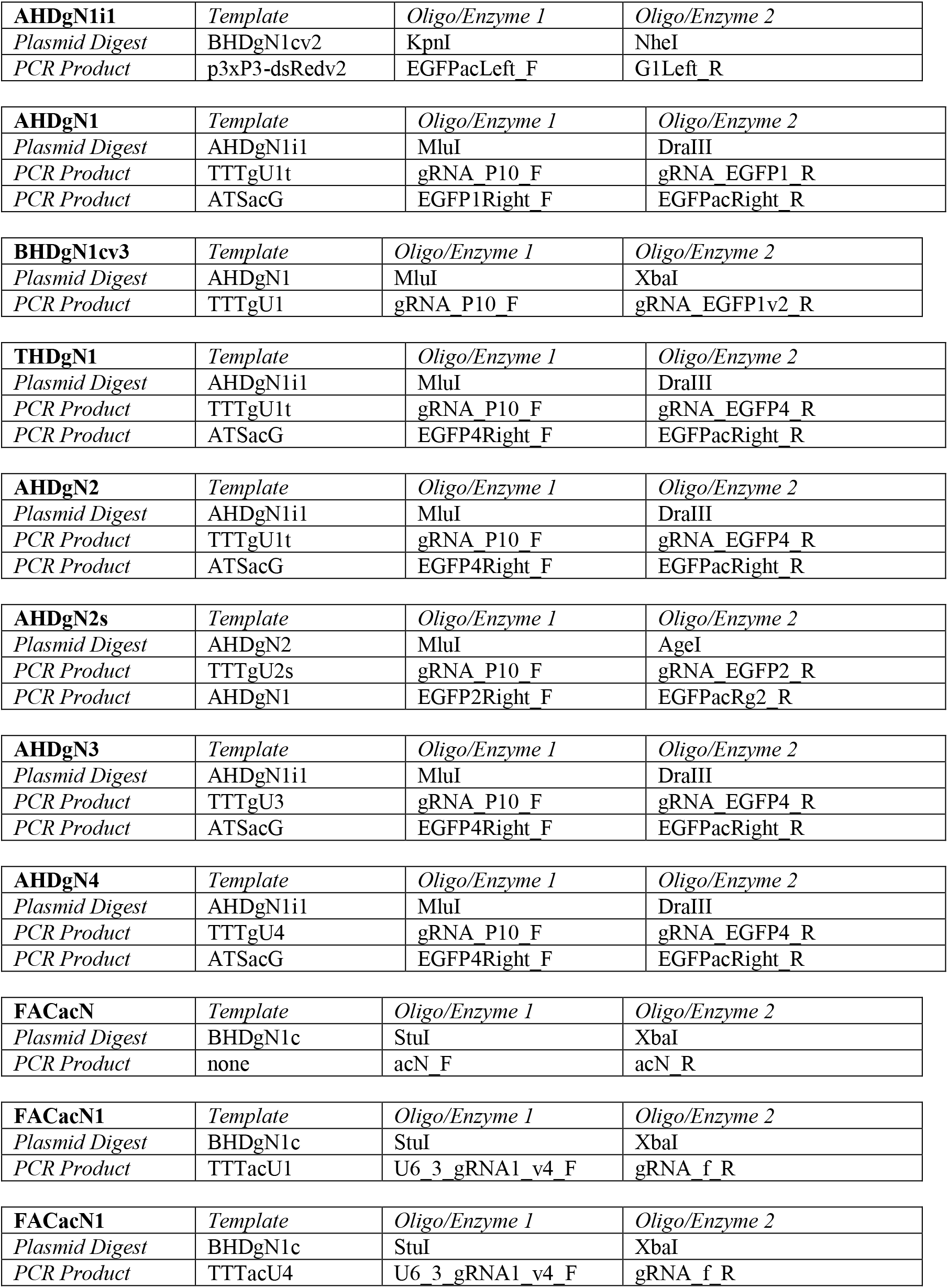

##### Construction primers

`Acg4_12_F: GGCAATATATAGGAATGCACGTTTTAGAGCTAGAAATAGCAAGTTAAA`

`Acg4_12_R: AACACTCGGTATAAATTGGTTTATGCACCAGCCGGGAATCG`

`Acg4_23_F: GCATAAACCAATTTATACCGAGTGTTTTAGAGCTAGAAATAGCAAGTTAAA`

`Acg4_23_R: AACTCCCCGCAAGTTCTGTCCCTTGCACCAGCCGGGAATCG`

`Acg4_34_F: GCAAGGGACAGAACTTGCGGGGAGTTTTAGAGCTAGAAATAGCAAGTTAAA`

`Acg4_34_R: GGTGGTCTCCGTTTTCCACTTGCACCAGCCGGGAATCG`

`Acg4_41_F: GTGCAAGTGGAAAACGGAGACCACCGTTTTAGAGCTAGAAATAGCAAGTTAAA`

`Acg4_41_R: AAAACGTGCATTCCTATATATTGCCTGCATCGGCCGGGAATCG`

`acgRNA_1_F: ACGTCGGCAATATATAGGAATGCACGTTTTAGAGCTAGAAATAGCAAGTTAAAATAAGG`

`acgRNA_2_R: AAAACGTGCATTCCTATATATTGCCGACGTTAAATTGAAAATAGGTCTATATATACG`

`acN_F: CAAACTCATCAATGTATCTTAACCGGTAGGAGCAAGCTGCCCGTGCCCTGGCCCACCCTC`

`acN_R: GAGGGTGGGCCAGGGCACGGGCAGCTTGCTCCTACCGGTTAAGATACATTGATGAGTTTG`

`AutoC_Left_F: ACATTATCGCGAGCCGACAGAAGAACGACCCGACAG`

`AutoC_Left_R: ATTAGATCCCGTACGACGTACCCATTGTTTGCTTTTAATCT`

`AutoC_Right_F: TATCTTAACCGGCGGAGGTGGCCATATCGCACTACA`

`AutoC_Right_R: GCAGAAGGCCCCTGACGACGGGCAAGGGAATTCAACA`

`b1v2_F: ATTTCGAGGTTAAAACGGTCGAAGCGCGGCCGCGGATCTAATTCAATTAGAGACTAATTC`

`b1v2_R: GAGTAGGAGCAATCACAGGTGAGCAAAAAAACGCGTGTTAACTCGAATCGCTATCCA`

`Cas9Nos3_R: TATCCACTTGTTTACTCTGACCAACT`

`Cas9Nos3c_F: TCTGCACCACCGGCTAGCTCCTTCCTGGCCCTTTTCGAG`

CFD_1_F: GTTTTAGAGCTAGAAATAGCAAGTTAAAATAAGG

CFD_1_R: GGCTATGCGTTGTTTGTTCTGC

CFD_2_F: AACAGTAGGCAGAACAAACAACGC

CFD35_2_R: CGACGTTAAATTGAAAATAGGTCTATATATACG

CFD5_2_R: TGCATCGGCCGGGAATCGA

`CFDg_1_F: GACCTGTTTTAGAGCTTTTTTGCCTACCTGGAGCCT`

`CFDg_2_R: CAGGTAGGCAAAAAAGCTCTAAAACAGGTCTTCTGCACCA`

`EGFP_c_F: AAACAATGGGTACGTCGTACGGGATCTAATTCAATTAGAGACTAA`

`EGFP_c_R: ATATGGCCACCTCCGCCGGTTAAGATACATTGATGAGTTTGG`

`EGFP1_gRNA_F: TATATATAGACCTATTTTCAATTTAACGTCGAAGTTCGAGGGCGACACC`

`EGFP1_gRNA_R: ATTTTAACTTGCTATTTCTAGCTCTAAAACGGGTGTCGCCCTCGAACTT`

`EGFP1Right_F: ATGCGTATGCATTCTAGACCCTGGTGAACCGCATCGAG`

`EGFP1t_gRNA_F: GCGGCCCGGGTTCGATTCCCGGCCGATGCAGAAGTTCGAGGGCGACACC`

`EGFP1tt_gRNA_F: GCGGCCCGGGTTCGATTCCCGGCCGATGCAGGTGGTGCAGATGAACTTCA`

`EGFP1t_gRNA_R: ATTTTAACTTGCTATTTCTAGCTCTAAAACTGAAGTTCATCTGCACCACC`

`EGFP2_gRNA_F: TATATATAGACCTATTTTCAATTTAACGTCGGGCACGGGCAGCTTGCCGG`

`EGFP2_gRNA_R: ATTTTAACTTGCTATTTCTAGCTCTAAAACCCGGCAAGCTGCCCGTGCCC`

`EGFP2Right_F: ATGCGTATGCATTCTAGATCAAGGAGGACGGCAACATCC`

EGFP4_12_F: GAAGTTCGAGGGCGACACCCGTTTTAGAGCTAGAAATAGCAAGTTAAA

EGFP4_12_R: AACACAAGCAGAAGAACGGCATCTGCACCAGCCGGGAATCG

EGFP4_23_F: GCAGATGCCGTTCTTCTGCTTGTGTTTTAGAGCTAGAAATAGCAAGTTAAA

EGFP4_23_R: AACGCTTGTGCCCCAGGATGTTGTGCACCAGCCGGGAATCG

EGFP4_34_F: GCACAACATCCTGGGGCACAAGCGTTTTAGAGCTAGAAATAGCAAGTTAAA

EGFP4_34_R: TGAAGTCGATGCCCTTCAGCTGCACCAGCCGGGAATCG

EGFP4_41_F: GTGCAGCTGAAGGGCATCGACTTCAGTTTTAGAGCTAGAAATAGCAAGTTAAA

EGFP4_41_R: AAAACGGGTGTCGCCCTCGAACTTCTGCATCGGCCGGGAATCG

`EGFP4Right_F: ATGCGTATGCATTCTAGAAGCAGAAGAACGGCATCAAGGTG`

`EGFPacLeft_F: ATTAACCAATTCTGAACATTATCGCCTAGGGTACCGACAGAAGAACGACCCGACAG`

`EGFPacLeft_R: GGCCAGGAAGGAGCTAGCCGGTGGTGCAGATGAACTTCA`

`EGFPacRg2_R: CAATTTTCCGTTGCACTTTTCGATTTCG`

`EGFPacRight_F: ATGCGTATGCATTCTAGAGCAAGCTGCCCGTGCCCT`

`EGFPacRight_R: TGATTGACGGAAGAGCCTCGAGCTGCACACACAGTGGACGGGCAAGGGAATTCAACATCC`

`EGFPc_U6_3_gRNA1_R: CGGGCAGCTTGCTCTAGAATGCATACGCATTAAGCGAACA`

`G1Left_R: GCGGCGTTTCTCGAAAAGGGCCAGGAAGGAGCTAGCTGTCGCCCTCGAACTTCAC`

`gRNA_EGFP1_R: GGTTCACCAGGGTCTAGAATGCATACGCATTAAGCGAACA`

`gRNA_EGFP1v2_R: GCCCTTCAGCTCGATGCGGTTCACCAGGGTCTAGAATGCATACGCATTAAGCGAACA`

`gRNA_EGFP2_R: CGTCCTCCTTGATCTAGAATGCATACGCATTAAGCGAACA`

`gRNA_EGFP4_R: CGTTCTTCTGCTTCTAGAATGCATACGCATTAAGCGAACA`

`gRNA_f_R: GAGGGTGGGCCAGGGCACGGGCAGCTTGCTCTAGAATGCATACGCATTAAGCGAACA`

`gRNA_P10_F: AGCTGGCTTGGATAGCGATTCGAGTTAACACGCGTTTTTTTGCTCACCTGTGATTGCTC`

`gRNAtRNA_F: ACATTATCGCGAGCCTTTTTTGCTCACCTGTGATTGCT`

`gRNAtRNA_R: CAGAAGGCCCCTGACATGCATACGCATTAAGCGAACA`

`pBB_1_R: GACCAAAATCCCTTAACGTGAGTT`

`pBB_2_F: GCGCGTAACTCACGTTAAGG`

`pBB2_c_F: ATTCCCTTGCCCGTCGTCAGGGGCCTTCTGCTTAGT`

`pBB2_c_R: GGTCGTTCTTCTGTCGGCTCGCGATAATGTTCAGAATTG`

`pBBtRNA_1_F: TTAATGCGTATGCATGTCAGGGGCCTTCTGCTTAGT`

`pBBtRNA_2_R: CAGGTGAGCAAAAAAGGCTCGCGATAATGTTCAGAATTG`

`U6_3_gRNA1_v4_F: GTCCAAACTCATCAATGTATCTTAACCGGTAGGCCTTTTTTTGCTCACCTGTGATTGCTC`

##### Sequencing primers

`AutoC_Left_S_F: AGCAGAGAAAAGTGTAGAGCACG`

`AutoC_Left_S_R: GTGCTGACCCACGATCCATTC`

`AutoC_Right_S_F: CCCCCTTCTGCACACCATACA`

`AutoC_Right_S_R: TACACCTCACACTACTCGGGC`

`AutoDLeft_S2_F: CTTACGCTGAAGCCATTTCAA`

`AutoDRight_S2_R: ATCTGGTTCTCACTTCCATTTAAAT`

EGFP_S_F: AGCGCACCATCTTCTTCAAGG

EGFP_S_R: AGTTGTACTCCAGCTTGTGCC

EGFP_S2_F: CCCTGAAGTTCATCTGCACCA

EGFP_S2_R: CTCCAGCAGGACCATGTGATC

IHD_S_F: GGGTTATTGTCTCATGAGCGG

IHD_S_R: TCTCGAAAATAATAAAGGGAAAATCAG

`pCFD5_S_R: ACGTCAACGGAAAACCATTGTCTA`

#### Generation of transgenic lines

Lines were transformed by Rainbow Transgenic Flies via injection of a donor plasmid (ATSacG, BHDgN1cv3, AHDgN1, THDgN1, AHDgN2, AHDgN2s, AHDgN3, AHDgN4, FACacN, FACacN1, FACacN4) into a *w*^*1118*^ (for ATSacG, FACacN, FACacN1, FACacN4) or into the ATSacG line (for the rest). Plasmid pHsp70-Cas9^34^ (provided by Melissa Harrison & Kate O’Connor-Giles & Jill Wildonger, Addgene plasmid #45945) was included in the injection as a source of Cas9 and plasmid BHDgg1c (for ATSacG, FACacN, FACacN1, and FACacN4), TTTgU1t (for BHDgN1cv3, AHDgN1, THDgN1), TTTgU2s (for AHDgN2s), or TTTgU4 (for AHDgN2, AHDgN3, AHDgN4) was included as a source of gRNA. Concentrations in the injection mix of donor, Cas9, and gRNA plasmids were approximately 500, 500, and 50 ng/µL, respectively in 10 mM Tris-HCl, 100 µM EDTA, pH 8.5 solution. Progeny of injected flies with dsRed fluorescent protein in the eyes, which usually indicated successful drive insertion, were crossed to each other for several generations to obtain homozygous stocks, with preference for flies with brighter eyes, which usually indicated that the individual was a drive homozygote. The stock was considered homozygous after sequencing confirmation. The split-CRISPR line with Cas9 driven by the *nanos* promoter and the driving component targeting *yellow* are detailed in a previous study^15^.

#### Fly rearing and phenotyping

Flies were reared at 25°C with a 14/10 hr day/night cycle. Fresh Bloomington Standard Medium was provided every two weeks. For phenotyping, flies were anesthetized with CO_2_ and examined with a stereo dissecting microscope. Red and green fluorescent eye phenotypes were scored using the NIGHTSEA system (SFA-GR and SFA-RB-GO). The different phenotypes and genotypes of our drive systems are summarized in the Supplemental Datasets, as are the calculations we used for determining drive performance parameters.

Experiments involving gene drive flies were carried out with Arthropod Containment Level 2 protocols at the Sarkaria Arthropod Research Laboratory at Cornell University, a quarantine facility constructed to containment standards developed by USDA APHIS. Additional safety protocols for insect handling were approved by the Institutional Biosafety Committee at Cornell University, further minimizing the risk of accidental release of transgenic flies. All drive flies also utilized our split-Cas9 system or synthetic target sites^15^, which should prevent the spread of the drive in the case of an accidental escape.

#### Genotyping

To obtain sequences of gRNA target sites, flies were frozen and homogenized in 30µL of 10 mM Tris-HCl pH 8, 1mM EDTA, 25 mM NaCl, and 200 µg/mL recombinant proteinase K (Thermo Scientific). The solution was incubated at 37°C for 30 min and then 95°C for 5 min. The mixture was used as the template for PCR to amplify the gRNA target sites. DNA was then was purified by gel extraction and Sanger sequenced. Sequences were analyzed with ApE software available at: http://biologylabs.utah.edu/jorgensen/wayned/ape.

#### Drive variants

In our model, we consider five types of homing gene drive systems:

1. Standard drive. The standard homing drive is a population modification system. Its primary drive mechanism occurs in germline cells during early meiosis. When it operates successfully, the drive allele replaces any wild type alleles in the germline. However, resistance alleles can also form, preventing the spread of the drive.
2. Population suppression drive. The drive increases in frequency in the same manner as the standard homing drive, and resistance alleles develop under the same circumstances. However, the drive targets a recessive female fertility gene, which disrupts the function of the gene with its presence. Resistance alleles can also disrupt the function of the target gene. Females with two disrupted copies of the gene are rendered sterile, while males are unaffected. Notably, unlike the standard homing drive, this drive does not carry any payload. The function of the drive is accomplished by suppressing the population. Such a drive was successful in laboratory populations of the mosquito *A. gambiae*^21^.
3. Haplolethal drive. This drive system is a modification of the standard homing drive system. It targets a gene that is critical to the viability of the individual. However, the drive contains a recoded portion of the gene that is immune to Cas9 cleavage, so the presence of the drive does not disrupt the function of the target. If any individual receives a resistance allele that disrupts the haplolethal target, that individual will not be viable, preventing such resistance alleles from entering the population. A haplolethal homing drive was successful in a laboratory population of the fruit fly *D. melanogaster*^16^.
4. Recessive lethal drive. This drive is similar to the haplolethal drive, but the target is recessive lethal. Only individuals carrying two resistance alleles that disrupt the target gene function will be nonviable. Thus, resistance alleles are removed from the population more slowly. However, this drive may be easier to engineer because the drive can provide rescue even in the presence of a resistance allele. It is also more tolerant of a high rate of embryo resistance allele formation because this would allow it to operate better as a toxin-antidote system^30,31^.
5. Gene disruption drive. The gene disruption homing drive is a population modification system that is similar to the suppression drive in that its presence disrupts the target gene, as do resistance alleles. However, individuals with two disrupted copies of this gene remain viable and fertile, though they suffer from a small additional fitness cost. The purpose of this drive is to remove the functionality of a particular gene from the population, which can provide benefits such as reduction of disease transmission^27,28^. An advantage of this drive is that there is no need for a recoded sequence. However, finding suitable targets for particular applications could potentially be difficult.

#### Computational model

We implemented each of the gene drive models using SLiM version 3.2.1^35^. SLiM is an individual-based, forward-time population genetic simulation framework. General parameters and ecology components are shared across all models.

Our model considers a single panmictic population of sexually reproducing diploid individuals with non-overlapping generations. The model differs from a standard Wright-Fisher type model in that population size is not regulated. Offspring are generated from random pairings throughout the population, with mate choice and female fecundity affected by genotype fitness. Fecundity is also multiplied by a factor representing the impact of the amount of crowding in the system: 10/(1+9N/K), where N is the total population and K is the carrying capacity. A number of offspring are then generated based on a binomial distribution with a maximum of 50 and *p* = fitness/25. This model produces logistic dynamics, while allowing the population size to fluctuate around the expected capacity. After pairings and offspring have been determined, the genotypes of the offspring are modified according to the genetic component of the model.

In one set of simulations, a small number of drive/wild-type heterozygous flies were introduced into a wild-type population of 100,000 at an initial frequency of 1%. The simulation was then conducted for 100 generations. In another set of simulations, a wild-type female was crossed to a drive/wild-type heterozygote male, and a configurable number of offspring were generated from that single pairing. The genotype of each offspring was recoded to estimate drive performance parameters. Drive conversion was equal to the fraction of wild-type alleles in the germline converted to drive alleles, and resistance allele formation rates also represented rates of conversion from wild-type alleles.

#### Genetic computational module

Except in the simple model described in the results, the flow for DNA modification events in our model is as follows: first, after generating an individual, both of the individual’s genes are subject to the formation of resistance alleles in a germline resistance function that retroactively describes changes that occurred in the germline cells of the parents; next, a homology-directed repair function determines whether an allele was converted to a drive allele; then, there is a second application of the germline resistance function, using a different resistance rate parameter; finally, an embryo resistance function determines whether Cas9 inherited from the mother forms any resistance alleles. Each of these functions, along with a Cas9 cutting function which is invoked by the other functions, is described below.

Germline resistance function:

This function runs on each chromosome, both before and after homology-directed repair. This function first determines a cut rate, and then passes that rate as an argument to a function that represents Cas9 possibly cutting and generating resistance alleles. The function only operates under the threshold conditions that the individual inherited a chromosome with at least one wild type locus from a parent that was a carrier for the drive (necessarily on the parent’s other chromosome). If these conditions are met, the rate of cutting is then determined. For a default rate of cutting, this function takes as an argument a global resistance rate as a parameter (see default parameters below).

However, one of the features of this model is the simulation of the possibility of simultaneous cleavage. When this feature of the model is activated, the cut rate is not simply the resistance rate parameter, but rather, the likelihood of cutting at each subphase is reduced to:

Subphase cut rate = 1 − (1 − resistance rate) ^ (1 / number of cut phases)

The subphase cut rate calculation is further modified when simulating Cas9 activity saturation. The per phase cut rate is then calculated as:

Subphase cut rate = 1 − (1 − resistance rate) ^ [Cas factor / (number of cut phases * number of gRNAs)]

where

Cas factor = Cas9 saturation parameter * number of gRNAs / (Cas9 saturation parameter + number of gRNAs − 1)

The final modification of this function is present when our model considers differing gRNA activity level at each different locus. In this case, the function generates a series of cut rates, rather than just one. This model has a global gRNA activity variation parameter. Based on this parameter, the range of gRNA activity multipliers at the target sites is then constructed as a list with a maximum of (1 + the parameter) and a minimum of (1 − the parameter), with the number of entries in the list equal to the number of gRNAs. The activity multiplier at each site steps down from the maximum to the minimum in linear steps. The nth cut rate is determined as follows:

Subphase cut rate = 1 − (1 − resistance rate) ^ (Nth Cas factor / (number of cut phases * number of gRNAs))

where

Nth Cas factor = Cas9 saturation parameter * number of gRNAs / (Cas9 saturation parameter + number of gRNAs − 1) * nth gRNA activity multiplier

After this function has determined the cut rate (or series of cut rates), it is passed as an argument to a Cas9 cutting function described below to determine if a resistance allele forms on the chromosome that the offspring is inheriting from the parent in question.

Embryo resistance function:

The function to determine resistance formation rates in the embryo is highly similar to the function that determines resistance rates in the germline. This function only proceeds when the threshold conditions of the mother being a carrier for the drive and the child having at least one wild type locus are met.

When these conditions are met, the calculations for per phase cut rate differ slightly from those in the germline function since it is affected by the number of copies of the drive present in the mother. The basic model is:

Subphase cut rate = 1 − (1 − resistance rate) ^ (mother drive count / number of cut phases)

The function also has one special case. For mothers that are drive/wild type, Cas9 activity has been determined to be higher than for mothers who are drive/resistance (see Supplemental Results). Thus, in the drive/wild-type case, the mother drive count variable is set to 1.83 for individuals that inherit a drive allele from the mother.

When modeling both saturation as well as variable gRNA activity level, the mother drive count has the same place in the resultant cut rate equation. The model including all of these factors calculates the cut rates as:

Subphase cut rate = 1 − (1 − resistance rate) ^ (mother drive count * Nth Cas factor / (number of cut phases * number of gRNAs))

where

Nth Cas factor = Cas9 saturation parameter * number of gRNAs / (Cas9 saturation parameter + number of gRNAs − 1) * nth gRNA activity multiplier

After the cut rate has been determined, it is passed as an argument to the Cas cut function which is run on both of the offspring’s chromosomes.

Cas cut function:

This function takes the cut rates determined by the above functions and modifies the chromosome. During each of the cuts, a random number between zero and one is checked against the cut rate for each wild type locus in the offspring’s chromosome. If, during any given cut phase, more than one site is cut, the left most locus is converted to a resistance allele that disrupts the function of the target gene, and the rest of the section between the two cuts and including the rightmost cut is marked with a placeholder that represents the absence of this section of DNA. If only one cut is made during a cut phase, the site is converted to either a resistance sequence that disrupts or preserves the function of the target gene at a specified rate.

Homology directed repair function:

This function determines whether a homing drive successfully copies itself onto the offspring’s chromosome. The function runs twice – once considering the offspring’s paternal chromosome and the father’s genome and once for the maternal chromosome. The function only runs when the threshold conditions are met of the offspring having a wild type locus on the chromosome it inherited from a parent who was a carrier for the drive. Before considering any special features, this function flows as follows:

First, a cut rate is passed to the function from a homing phase cut rate default parameter. Each wild type locus on the chromosome is checked against that cut rate. If any cuts are made, an additional check is made against a baseline homing success rate parameter. If homing succeeds, the chromosome is converted to a drive chromosome. If Cas9 cuts, but homing fails, end-joining repair occurs. In this case, as in the Cas cut function, if multiple loci were cut, the span of DNA is marked as missing, and the first locus is marked as a function disrupting resistance allele. If only one cut is made, either type of resistance sequence can form at the site.

When modeling gRNA saturation, the cut rate is altered to include a factor related to the Cas9 saturation factor as well as the number of gRNAs, much like the cut rate is altered in resistance formation. When simulating gRNA saturation, the homology-directed repair cut rate is:

Cut rate = 1 − (1 − homing phase cut rate parameter) ^ (Cas factor / number of gRNAs)

where

Cas factor = Cas9 saturation parameter * number of gRNAs / (Cas9 saturation parameter + number of gRNAs − 1)

Just as resistance allele formation can be toggled to vary at each different locus, the cut rates in the homology-directed repair phase can also be modified to reflect variable gRNA activity level at each locus. When toggled on, the cut rate is as follows:

Per phase cut rate = 1 − (1 − homing phase cut rate parameter) ^ (Nth Cas factor / number of gRNAs)

where

Nth Cas factor = Cas9 saturation parameter * number of gRNAs / (Cas9 saturation parameter + number of gRNAs − 1) * nth gRNA activity multiplier

When the entire target area of the drive is not cut out, the excess DNA between the outer target sites and the closest sites that were cut results in lower probability of homing occurring successfully (referred to as a repair fidelity penalty). The next toggleable feature of the model is the simulation of these cut offset effects. After it is determined at which loci Cas9 cuts, the model determines how close the left and right cut edges are to the leftmost and rightmost target loci (by considering the index of the cut and also accounting for the fact that segments of the genome may have previously been excised due to simultaneous cutting during resistance formation). The offsets from the actual leftmost and rightmost sites modify the rate of homing success as follows:

Homing success rate = baseline success parameter * (1- homing edge effect parameter * left offset) * (1- homing edge effect parameter * right offset)

For example, consider a genome with 10 target loci. Cas9 has made cuts at sites 3, 4, 6, and 7, and DNA spanning sites 8, 9, and 10 was previously removed due to simultaneous cutting at sites 8 and 10 during resistance formation. Thus, the left side offset is 2 (site three is two away from the actual leftmost locus) and the right offset is 1 (site 7 is the rightmost cut target site, but there is only one locus with any genetic material present to the right of it).

The final toggleable modification in this function is ability to simulate incomplete homology-directed repair. When this feature is enabled, in cases that Cas has cut, but when the drive has failed to successfully home, there is a chance that the drive will experience incomplete homology-directed repair failure. This will convert the allele into a full resistance allele that disrupts the function of the target gene. The odds of this occurring are related to the cut offsets, as described above. The incomplete homology-directed repair rate is given by:

Incomplete homology-directed repair rate = 1 - base total failure avoidance rate parameter * (1 − 0.1 * left offset) * (1 − 0.1 * right offset)

If incomplete homology-directed repair does occur, then drives with haplolethal or recessive lethal target sites (those with recoded versions of these genes in the drive) have an additional chance of the allele being converted to a full resistance allele that preserves the function of the target gene. The rate of this occurring is:

Rate = rate parameter * (1+ right offset − left offset)

The asymmetry in this function between the right and left offsets is because the recoded region is assumed to be on the left end of the homing drive.

#### Summary of default model parameters

drive homozygote fitness value: 0.9

drive heterozygote fitness value: 0.949

additional gene disruption drive fitness multiplier for individuals with two copies of the drive and/or resistance alleles that disrupt the function of the target gene: 0.95

early germline resistance formation phase cleavage rate: 0.02

late germline resistance formation phase cleavage rate: 0.9

homology-directed repair phase cleavage rate: 0.98

baseline rate at which homology-directed repair (as opposed to end-joining repair) occurs in the homology-directed repair phase after cleavage: 0.95

embryo resistance formation phase cleavage rate: 0.05

chance to form a function preserving resistance allele at a cleavage site: 0.1

number of subphases in resistance formation phases: 3

gRNA activity variation level: 0.2

repair fidelity penalty per step length lacking homology to the drive: 0.055 Cas9 activity saturation level: 1.5

incomplete homology-directed repair baseline rate: 0.1

formation of a complete resistance allele that preserves the function of the target gene if a recoded region is present (haplolethal and recessive lethal drives) and incomplete homology-directed repair takes place rate: 0.001

carrying capacity of the environment: 100,000

drive/wild-type heterozygote release size: 100

#### Model and data availability

All SLiM configuration files for the implementation of the different models and all simulation data are available on GitHub (https://github.com/MesserLab/Homing_Mechanisms_with_multiplexed_gRNAs).

## SUPPLEMENTAL RESULTS

**Figure S1.**
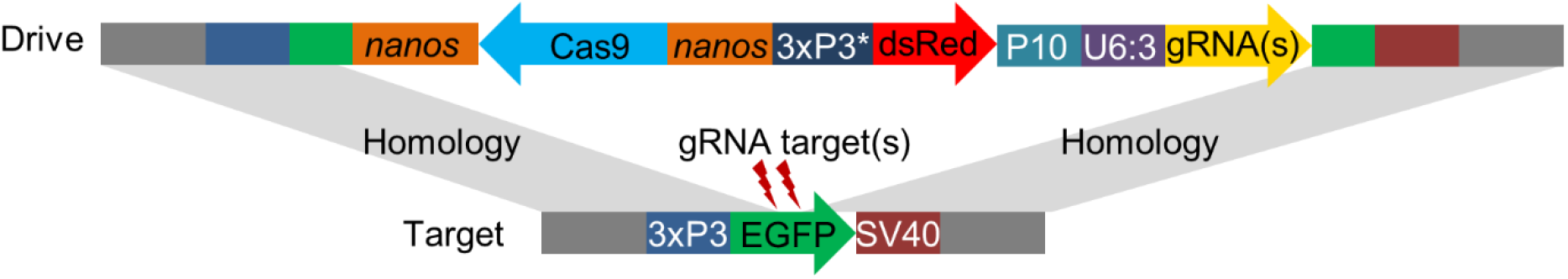
Synthetic target site drive schematic diagram. The synthetic target site drive constructs contain Cas9 with the germline *nanos* promoter and 3’UTR, a dsRed marker with a slightly recoded (*) 3xP3 promoter and P10 3’UTR, and U6:3 promoter driving one or more tRNA-linked gRNAs that target EGFP. The homology arms include the EGFP target sequence around the outer cut sites together with the 3xP3 promoter and SV40 3’UTR regions.

### Cut site sequence analysiss

Progeny of drive/wild-type heterozygote females that contained the drive but did not have EGFP phenotype were sequenced to determine the pattern of resistance alleles at each gRNA target site.

**Table S1.**

| **gRNAs present in mother** | **Cut site 1** | **Cut site 2** | **Cut site 3** | **Cut site 4** | **# of sequences with pattern** |
| --- | --- | --- | --- | --- | --- |
| 1,2,3,4 | R | mosaic | WT | WT | 2 |
| 1,2,3,4 | R | WT | mosaic | WT | 3 |
| 1,2,3,4 | R | WT | R | WT | 3 |
| 1,2,3,4 | R | WT | WT | mosaic | 1 |
| 1,2,3,4 | R | WT | WT | WT | 5 |
| 1,2,3,4 | R- | - | - | -R | 1 |
| 1,2,3,4 | R- | - | -R | WT | 9 |
| 1,2,3,4 | WT | R | WT | WT | 1 |
| 1,2,3,4 | WT | WT | mosaic | WT | 1 |
| 1,3,4 | R | WT | mosaic | WT | 1 |
| 1,3,4 | R | WT | R | mosaic | 1 |
| 1,3,4 | R | WT | R | WT | 1 |
| 1,3,4 | R | WT | WT | WT | 3 |
| 1,3,4 | R- | - | -R | WT | 8 |
| 1,3,4 | R- | -R | mosaic | WT | 1 |
| 1,4 | R | WT | WT | mosaic | 1 |
| 1,4 | R | WT | WT | WT | 6 |
| 1,4 | R- | - | - | -R | 1 |

WT = wild-type sequence
R = resistance sequence
“-“ = large deletion between cut sites

#### Additional timing components of the model

In our model, resistance alleles are first formed in the early germline. Each wild-type gRNA target site has a 2% probability of being cut. All cuts at this stage undergo end-joining repair, resulting in the formation of resistance alleles. Next is the homology-directed repair phase. Each remaining wild-type site has a 98% probability of being cut. If any sites are cut, homology-directed repair occurs in 95% of cases, resulting in conversion of the entire allele to a drive allele. Otherwise, end-joining repair and formation of resistance alleles takes place as described above. These parameters produce a drive with a conversion efficiency of 91% and inheritance of 96% for one gRNA, which is similar to existing *Anopheles* homing drives^18,19,21,25^. Finally, since it appears that most wild-type alleles are converted to resistance alleles in the germline^12,13,15^, we add another late germline resistance allele formation phase with a high cut rate of 90%, resulting in few wild-type alleles remaining. If at any stage multiple sites are cut, the region between them is deleted (preventing future cleavage of deleted gRNA target sites). Overall, in this model, adding additional gRNAs is beneficial, but a maximum efficiency that is determined by the success rate of homology-directed repair is eventually reached (Figure 4).

In addition to those formed in the germline, resistance alleles also form in the early embryo due to maternal deposition of Cas9 and gRNA. Thus, any wild-type alleles obtained from the female or male parents can be cut if the female parent has at least one drive allele, regardless of whether a drive allele was actually inherited by the embryo. If the female has two drive alleles, the enzymatic activity of Cas9 is doubled, which somewhat increases the cleavage rate (see Methods). If the female has a drive allele and the other allele has at least one wild-type site, then it is likely that drive conversion occurs, resulting in increased deposition of Cas9 and gRNA into most embryos receiving the drive allele and thus, increased cleavage. To determine the rate of enzymatic activity in these embryos, we analyzed embryo resistance allele formation rates in the progeny of female drive/wild-type heterozygotes for drives targeting *yellow*^12^, *white*^13^, *cinnabar*^13^, and EGFP^15^ (also including the lines in this study) in the *w*^*1118*^ background, plus additional drives targeting *yellow* that were introduced into the Canton-S^12^, Global Diversity Line^12^, and *Drosophila* Genetic Reference Panel^14^ backgrounds. We found that an enzymatic activity level of 1.83 minimized the sum of squares for the difference between the predicted resistance allele formation in individuals inheriting a drive from a female heterozygote and the actual values. Such predictions were based on the embryo resistance rates in individuals not inheriting a drive allele, which were considered to have a Cas9/gRNA enzymatic activity level of 1. We therefore use this value in our model. We also use a low embryo cut rate of 5%, which appears to be similar to the rate in *Anopheles gambiae* drives using the *zpg* promoter^21,25^.

Additionally, these processes do not necessarily resolve themselves instantaneously. While the window for homology-directed repair is likely narrow, resistance allele formation can occur over an extended period of time either before or after this window. We therefore break up each resistance allele formation phase in both the germline and the embryo into several subphases, with cut rates adjusted such that the final probability of cutting a particular target site after all subphases are completed is equal to the originally specified cut rate parameter for the entire phase. Greater numbers of subphases result in less simultaneous deletion of target sites. This can be important for resistance allele formation, but it has a negligible effect on drive conversion efficiency. The effects of this mainly come into play in later generations by controlling the rate that segments are deleted during the formation of resistance alleles. A two-gRNA drive cuts its target sites simultaneously in one third of cases that formed resistance alleles^13^. We therefore move forward with three subphases in our model for all resistance allele formation phases.

#### Model with repair fidelity

To model repair fidelity, we modify the probability of successful drive conversion in the homology-directed repair phase. We assume that the reduction in the success rate is proportional to the length of the DNA segment lacking homology and to a repair fidelity penalty parameter. We further assume that gRNA cut sites are evenly spaced, so we measure length in terms of number of cut site intervals, or “steps” between the outer sites and the closest cleavage site. Penalties from left and right homology mismatches are assumed to be multiplicative (see Methods for details). As the penalty increases, additional gRNAs do not contribute substantially to drive conversion efficiency and overall drive conversion efficiency is reduced (Figure S2). However, though efficiency does not increase, neither is it reduced by a high number of gRNAs (Figure 4), since the cut rate at the outermost gRNAs remains constant.

**Figure S2.**
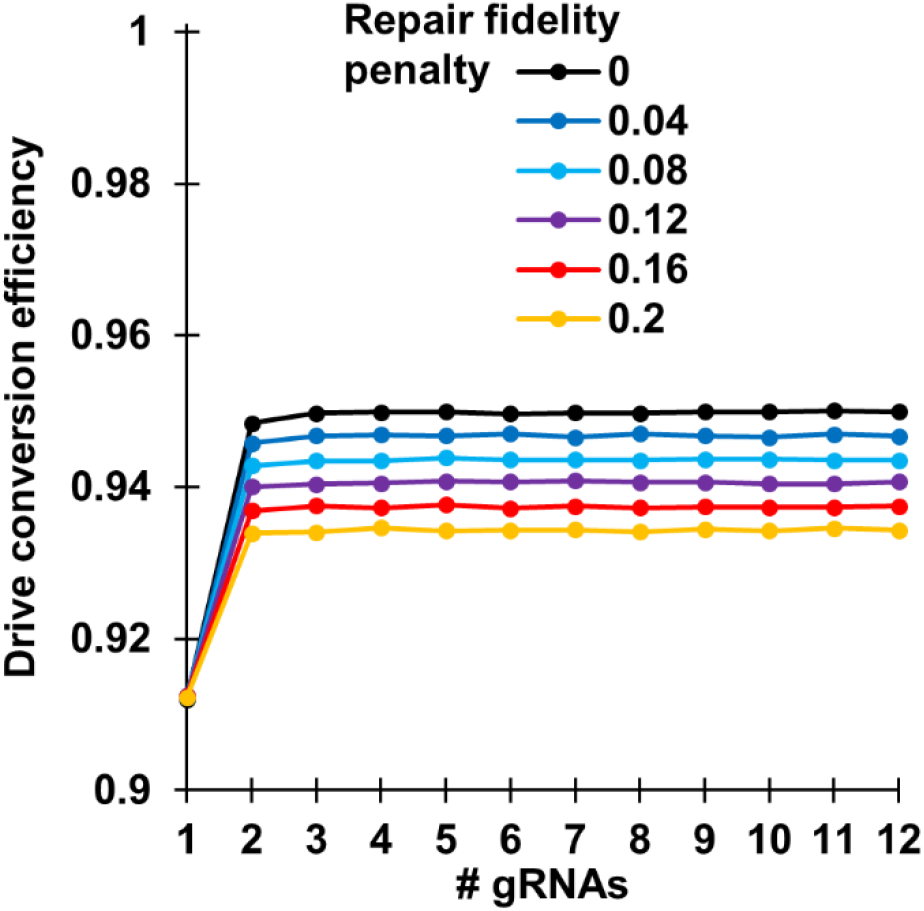
Effects of repair fidelity on drive performance. Five million offspring were generated from crosses between drive/wild-type heterozygotes and wild-type individuals for each number of gRNAs and repair fidelity penalty rate. The model including timing and repair fidelity components, but not other model features. The rate at which wild-type alleles are converted to drive alleles in the germline of drive/wild-type individuals is shown.

We estimated the value of the repair fidelity penalty parameter based on our experimental crosses. Selecting the female *Drosophila* drive/wild-type heterozygotes that have more similar performance to the highly efficient mosquito drives, we note that drive conversion in the drive with a poor right homology arm was 84% the value of the one-gRNA drive with ideal homology arms (Data S3). The right homology arm mismatch was equivalent to our four-gRNA drive if only the first gRNA cut, thus creating a right arm mismatch of three gRNA “steps” in the drive with a poor right homology arm. We therefore estimate the level of mismatch repair fidelity parameter to be a 5.5% efficiency reduction per gRNA step.

#### Model with and Cas9 activity saturation

To model Cas9 activity saturation, we simply reduce the cut rate per gRNA, with the overall cut rate (total Cas9 enzymatic activity) of all drives increasing asymptotically to a specified maximum Cas9 activity level, as specified in the methods. Since overall cleavage rates plateau, additional gRNAs beyond the first several do not substantially increase the rate of drive conversion (Figure 4, Figure S3).

**Figure S3.**
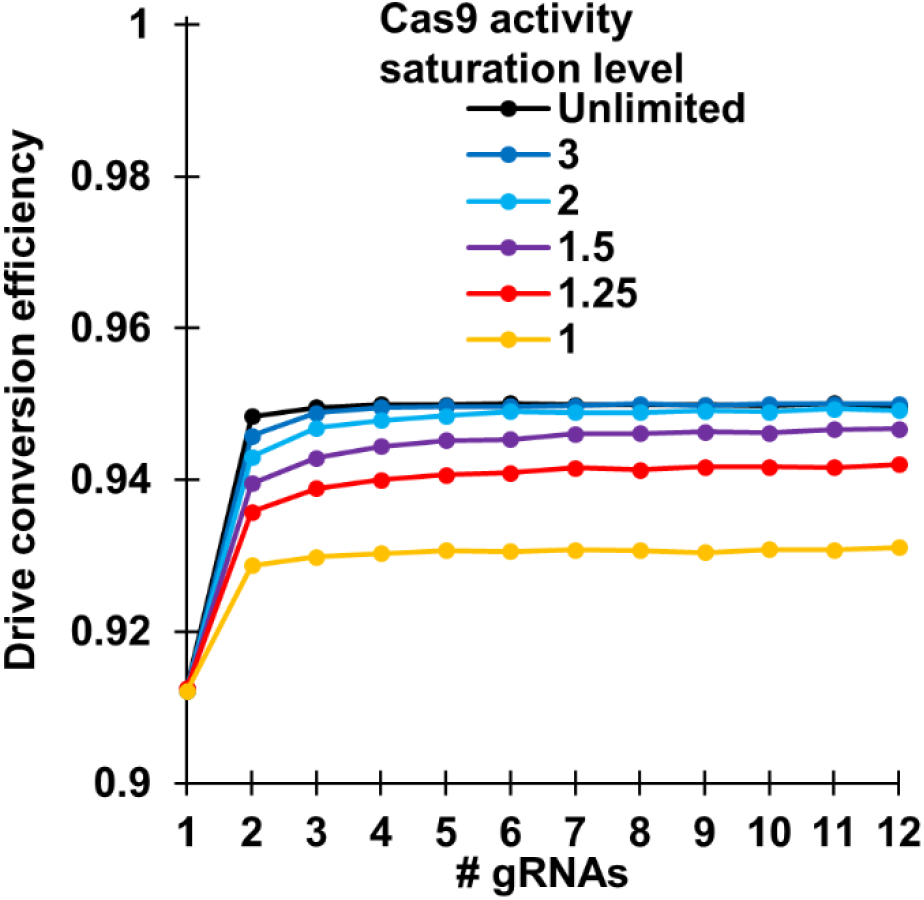
Effects of Cas9 activity saturation on drive performance. Five million offspring were generated from crosses between drive/wild-type heterozygotes and wild-type individuals for each number of gRNAs and Cas9 activity saturation level. The model including timing and Cas9 activity saturation components, but not other model features. The rate at which wild-type alleles are converted to drive alleles in the germline of drive/wild-type individuals is shown.

To estimate the Cas9 activity saturation parameter, we have two methods, each based on examining embryo resistance. First, using the split-*yellow* drives (Data S4) and taking into account copying of the gRNA (making its quantity equal to 1.83 times the quantity of other individual gRNAs for embryo resistance in individuals inheriting the drive, see timing section), we obtain values of 1.5 and 3.7 comparing the one-gRNA split-Cas9 and four-gRNA split-Cas9 alleles respectively to the baseline provided by the split-Cas9 without any gRNAs. However, the *yellow* gRNA is expressed at a different genomic location and without the tRNA system of the other gRNAs, which potentially accounts for the wide discrepancy between the two values. Another way to assess this parameter is to compare the embryo resistance of the one-gRNA drives with a tRNA to the embryo resistance rate of two-gRNA drives. This is because the second gRNA in each of these drives provides negligible cutting compared to the first in the embryo (Table S1). In our analysis, we focus on embryo resistance in flies that do not inherit the drive allele because Cas9 activity is overall lower, allowing a reduction in cleavage rate to be detected more easily despite the lower number of counts for these groups. This yields Cas9 maximum activity parameters of 1.6, 1.2, 1.9, and 1.5 when comparing the standard one-gRNA drive to the further spaced two-gRNA drive, the standard one-gRNA drive to the close spaced two-gRNA drive, the one-gRNA drive with poor right end homology to the further spaced two-gRNA drive, and the one-gRNA drive with poor right end homology to the close spaced two-gRNA drive, respectively. We therefore proceed with an estimate of 1.5 for the maximum Cas9 activity saturation level parameter.

#### Model with varying gRNA activity level

As indicated in our experiments, the relative activity levels of gRNAs can vary considerably, even if all are expressed together at presumably the same levels. We thus added a simplified version of gRNA activity variance to our model. This is based around a parameter that modifies the enzymatic activity level of each gRNA (see Methods). The left gRNA has its activity increased by a gRNA activity variance parameter, and the right gRNA has its activity decreased by this amount. Middle gRNAs are evenly “stepped down” in activity from left to right. With increasing gRNA activity variance, cleavage at gRNA sites near the right end is reduced, resulting in lower drive conversion due to the repair fidelity penalty (Figure S4). gRNAs with particularly low activity are unlikely to be used in drives designed for deployment in natural populations, but it is likely that there would still be some variance in gRNA activity. We thus selected 0.2 as the default parameter for gRNA activity variance, which has a small negative effect on drive conversion efficiency (Figure 4).

**Figure S4.**
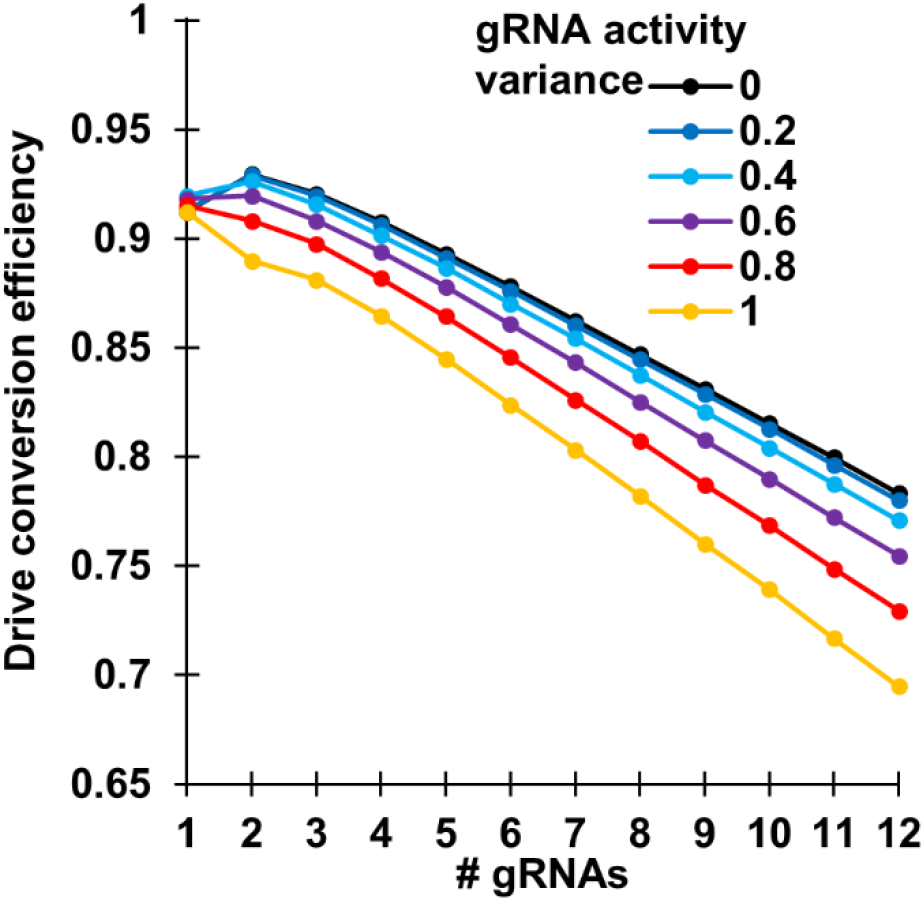
Effects of gRNA activity variance on drive performance. Five million offspring were generated from crosses between drive/wild-type heterozygotes and wild-type individuals for each number of gRNAs and gRNA activity variance level using the full model. The rate at which wild-type alleles are converted to drive alleles in the germline of drive/wild-type individuals is shown.

#### Incomplete homology-directed repair

Previous work has shown that some resistance alleles can be formed when homology-directed repair is interrupted, leaving a short sequence from the drive allele that is sufficient to disrupt any target gene and prevent future Cas9 cleavage. Based on sequencing, approximately 3% of resistance alleles for a drive targeting *yellow*^12^ and 7% for a drive targeting *white*^13^ were alleles formed by incomplete homology-directed repair. We thus model that in the homology-directed repair phase, if drive conversion does not occur, there is a 5% chance that incomplete homology-directed repair occurs, which converts all target sites into resistance alleles, even where cleavage did not take place. This chance is slightly increased if there is mismatch between ends in the same manner that drive conversion is decreased due to reduced repair fidelity (see Methods). This is because homology-directed repair may start at one chromosomal end with good homology, but it may fail at the other end where poor homology makes the process more difficult, resulting in incomplete homology-directed repair before end-joining mechanisms finish repair of the DNA. In most cases, incomplete homology-directed repair does not substantially increase the number of resistance alleles formed. However, in some cases, it can have a substantial effect due to the specific type of resistance alleles formed.

One strategy for designing efficient population modification drives. By choosing such targets, resistance alleles that disrupt the function of the target gene do not enter the population or carry high fitness costs. However, in such drives, incomplete homology-directed repair could result in copying of the recoded portion of these drives, but not the desired payload. This results in the formation of a complete resistance allele that preserves the function of the target gene. In our model, we include a parameter representing the chance that this occurs, given that incomplete homology-directed repair occurs. It is likely to be a rare phenomenon, but with even low rates, the formation rate of resistance alleles that preserve the function of the target gene can substantially increase in drives with several gRNAs (Figure 5). Without any information to estimate this parameter, we assume a default value of 0.1%. This places the rate on the order of the chance that a payload gene would be inactivated by mutations that form during homology-directed repair (estimated as approximately one in ten thousand per instance of homology-directed repair of the drive^36^).

#### Effect of cleavage rates on the performance of multiple gRNA homing drives

Our analysis used parameters inspired by highly efficient gene drives in *Anopheles*, but less efficient drives could still succeed in modifying or suppressing populations. Such drives may have a different optimal number of gRNAs. To investigate this, we examined a drive with similar performance to our synthetic target site drives in *D. melanogaster* constructed in this study, albeit with a reduced rate of early embryo resistance allele formation that would be necessary for the drives to be successful in at least some circumstances. Specifically, the early germline resistance allele formation phase cleavage was increased from our default of 2% to 5%. The homology-directed repair phase cleavage rate was reduced from 98% to 92%, and the embryo resistance allele formation phase cleave rate was increased from 5% to 10%. With these parameters, the performance of population modification drives was moderately worse, as expected (Figure S5). The optimal number of gRNAs for most of the drives was increased from three to four, and the negative effects of a high number of gRNAs were more pronounced. The success rates of suppression drives were more drastically impacted (Figure 7).

**Figure S5.**
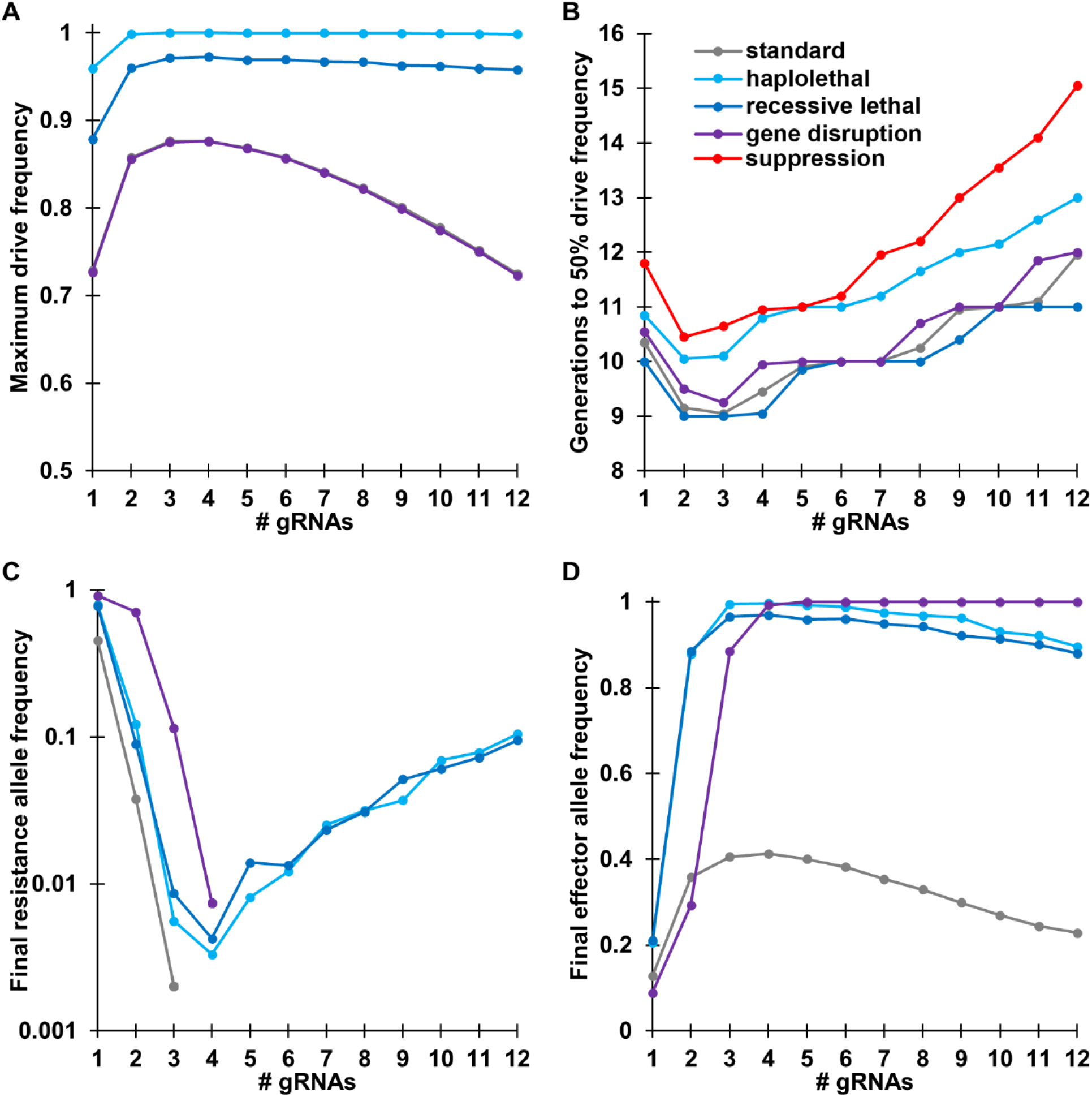
Comparison of performance parameters for different types of homing drives with lower cleavage efficiencies. Drive/wild-type heterozygotes were released into a population of 100,000 individuals at an initial frequency of 1%. The simulation was then conducted for 100 generations using the full model, but with reduced drive efficiency compared to the default parameters. The displayed results are the average from 20 simulations for each type of drive and number of gRNAs (**A**) The maximum drive allele frequency reached at any time in the simulations. Note that the standard drive and gene disruption drive values are highly similar. (**B**) The number of generations needed for the drive to reach at least 50% total allele frequency. Note that the suppression drive is only shown in (**B**). (**C**) The final frequency of resistance alleles after 100 generations. The displayed values are only for resistance alleles that preserve the function of the target gene. No resistance alleles were present in the standard drive and gene disruption drive when at least four gRNAs were present. (**D**) The final effector frequency present in the population after 100 generations. This was the drive allele only for most drive types, but for the gene disruption drive, it includes resistance alleles that disrupt the function of the target gene.

